# A network of small RNAs regulates sporulation initiation in *C. difficile*

**DOI:** 10.1101/2022.10.17.512509

**Authors:** Manuela Fuchs, Vanessa Lamm-Schmidt, Tina Lenče, Johannes Sulzer, Arne Bublitz, Milan Gerovac, Till Strowig, Franziska Faber

## Abstract

The obligate anaerobic, enteric pathogen *Clostridioides difficile* persists in the intestinal tract by forming antibiotic resistant endospores that contribute to relapsing and recurrent infections. Despite the importance of sporulation for *C. difficile* pathogenesis, environmental cues, and molecular mechanisms regulating sporulation initiation remain ill defined. Here, using RIL-seq to capture the Hfq-dependent RNA-RNA interactome, we discovered a network of small RNAs that bind to mRNAs encoding sporulation-related genes. We show that two of these small RNAs, SpoX and SpoY, regulate translation of the master regulator of sporulation, Spo0A, in an opposing manner, which ultimately leads to altered sporulation rates. Infection of antibiotic-treated mice with SpoX and SpoY deletion mutants revealed a global effect on gut colonization and intestinal sporulation. Our work uncovers an elaborate RNA-RNA interactome controlling the physiology and virulence of *C. difficile* and identifies a complex post-transcriptional layer in the regulation of spore formation in this important human pathogen.

## INTRODUCTION

Since its discovery as a causative agent of antibiotic-associated pseudomembranous colitis, *Clostridioides difficile* (*C. difficile*) has emerged as the leading cause of nosocomial antibiotic-associated disease in the developed world^1–3^. Several virulence traits contribute to disease severity of *C. difficile* infections (CDI), including exotoxin production and spore formation^4^. In particular, spores are a key element in host transmission and disease recurrence, due to their resistance to conventional antibiotics, disinfectants and other environmental stressors^5–7^. Hence, understanding the environmental signals and molecular mechanisms that control spore formation in this important human pathogen is essential for the development of alternative treatment options.

Spore formation has been studied extensively in a variety of sporulating bacteria and represents an energetically costly, morphogenic process that is irreversible beyond a certain point in spore development^8,9^. In particular, sporulation initiation is tightly controlled through the integration of environmental and nutritional signals that mediate the post-translational activation of the master regulator of sporulation, Spo0A^10,11^. However, *C. difficile* lacks many of the known conserved regulatory mechanisms that activate Spo0A, rendering sporulation initiation a poorly understood process in this gram-positive pathogen^12^. Once activated, phosphorylated Spo0A-P acts as a transcriptional regulator that induces the expression of a set of early sporulation genes. This ultimately leads to the hierarchical activation of four compartment-specific sigma factors - σ^E^/σ^K^ in the mother cell and σ^F^/σ^G^ in the forespore - and culminates in the formation of a metabolically dormant spore^13^.

Most recently, post-transcriptional regulation mediated by the RNA binding protein (RBP) Hfq has been implicated in modulating sporulation in *C. difficile*^14,15^. Boudry *et al*. demonstrated that depletion of Hfq leads to the upregulation of several sporulation related genes as well as an increased sporulation rate^14,15^. Hfq is known for its ability to facilitate base-pairing between small regulatory RNAs (sRNAs) and their target mRNAs, leading to altered translational efficiency and mRNA stability^16^. Similar to its extensively studied gram-negative counterparts, Hfq immunoprecipitation followed by sequencing of bound RNA species (RIP-seq) in *C. difficile* uncovered a vast number of sRNAs and mRNAs bound by Hfq^17,18^. Furthermore, several sRNAs, not only in *C. difficile* but also in other spore-forming Firmicutes, have been associated with the sporulation process, mostly through RNA-seq and microarray based expression profiles^14,19–21^. However, only a few of these sRNAs have been functionally described. In *C. difficile*, sRNA RCd1, that inhibits the production of the late mother cell-specific sigma factor σ^K^, remains the only sporulation associated sRNA characterized to this date^17^, revealing a paucity of knowledge that clearly warrants further investigation.

Global approaches such as RIP-seq are powerful tools in discovering RBP-bound sRNAs or mRNAs^18,22^. However, they rely on additional experimental and computational assays to identify directly interacting sRNA-target pairs^23^. Melamed and colleagues circumvented this difficulty by introducing RIL-seq (RNA interaction by ligation and sequencing) to the field of bacterial RNA-biology^24,25^. Similarly to CLASH and hiCLIP, RIL-seq relies on ligation of RBP-bound RNA pairs and thereby directly captures and identifies interaction partners^26,27^.

In the present study, we applied Hfq RIL-seq to *C. difficile*, which led to the discovery of an extensive Hfq-mediated sRNA-target network. Among the identified sRNA-mRNA interactions were several sRNAs bound to the *spo0A* mRNA, encoding the master regulator of sporulation. We show that two of these sRNAs, SpoY and SpoX, regulate *spo0A* translation in an opposite manner *in vivo*, resulting in altered sporulation rates. Furthermore, SpoY and SpoX deletion significantly impacts *C. difficile* gut colonization and spore burden in a mouse model of *C. difficile* infection. Overall, we provide the first example of sRNAs regulating sporulation initiation by finetuning *spo0A* translation, which adds a new layer of post-transcriptional regulation to the complex process of sporulation initiation in this important human pathogen.

## RESULTS

### Hfq is a global RNA binding protein that mediates sRNA-mRNA interactions in C. difficile

To better understand the impact of post-transcriptional regulation on sporulation, we performed Hfq RIL-seq in *C. difficile* in sporulating conditions^25^. *C. difficile* 630 cells expressing a chromosomally FLAG-tagged Hfq variant (Hfq-FLAG, n=4) were harvested during the transition phase, when *C. difficile* shifts to a non-growing state, accompanied by sporulation to ensure survival in nutrient limiting conditions^28,29^. Harvested cells were UV-crosslinked to stabilize *in vivo* protein-RNA interactions, followed by cell lysis and Hfq co-immunoprecipitation. Identification of Hfq-associated RNA-RNA interaction partners was achieved by ligation of Hfq-bound RNA pairs (“chimeras”), followed by RNA purification, sequencing, and computational analysis (Figure 1A). *C. difficile* 630 expressing native Hfq (WT) served as a control and was treated similarly (n=4). Analysis of the RIL-seq data revealed a high number of Hfq-bound single and chimeric fragments with a considerable enrichment of chimeric reads in the Hfq-FLAG strain, when compared to the WT (Figure 1B). The list of chimeras was manually curated and further reduced to statistically relevant interactions (Odds ratio ≥1 and p-value <0.05) that are represented by at least 25 chimeric fragments^25^. All remaining interactions are listed in Supplementary Table 2&3. The resulting RIL-seq network is publicly available and explorable in an RNA-RNA interactome browser (available upon publication, Supplementary Figure 1). In accordance with existing *E. coli* and *S. enterica* RIL-seq data, most chimeras (n=67%) consisted of mRNA-sRNA interactions, with mRNAs (5’UTR, CDS or 3’UTR) at position 1 (RNA1/5’end) and sRNAs at position 2 (RNA2/3’end), as shown in Figure 1C&D and Supplementary Figure 1B. Interestingly, sRNAs generally showed a clear preference for either position 1 or 2. For example, chimeric reads mapping to nc083 were consistently found at position 1, while chimeric reads mapping to nc159 were mostly located at position 2 within a chimera (Supplementary Figure 1B). However, of all chimeric fragments mapping to sRNAs more than 90% mapped to RNA2 (Figure 1D). This position bias reflects the mechanism by which most sRNAs bind Hfq in gram-negative species. Interactions generally occur between the proximal face of Hfq and the distinct intrinsic terminator and poly-U tail that characterizes most sRNAs, ultimately rendering the sRNA 3’end inaccessible to proximity ligation (Figure 1A) ^30^. Accordingly, our data imply that *C. difficile* Hfq might employ binding mechanisms similar to those described for *S. enterica* and *E. coli* in facilitating sRNA target interactions^30^. While the majority of sRNAs were found ligated to CDSs (56%), a surprisingly high number (33%) interacted with mRNA 3’UTRs (Figure 1C). Recently published *C. difficile* Hfq RIP-seq data revealed similar distributions of Hfq-bound RNA species, however, RIP-seq does not allow identification of direct interaction partners^18^. In contrast, sRNA-mRNA chimeras in *E. coli* and *S. enterica* were clearly dominated by sRNAs interacting with CDSs or 5’UTRs, while sRNA-3’UTR ligations were barely found^24,31^. Although bacterial 5’UTRs have long been described as the prototypical target of sRNA-mediated post-transcriptional regulation, there are examples of sRNAs targeting mRNA 3’UTRs, including sRNA Spot42 targeting the 310-nt long *hilD* 3’UTR, a transcriptional regulator of virulence in *S. enterica*^32–34^. Indeed, research on *S. aureus* suggests that long 3’UTRs in particular might be an underrated source of regulatory elements that impact transcript stability and translation^32,35^. Considering that in *C. difficile* 42% of all annotated 3’UTRs are longer than 100 nt, they might constitute a source of regulatory elements targeted by sRNAs^18^.

**Figure 1:**
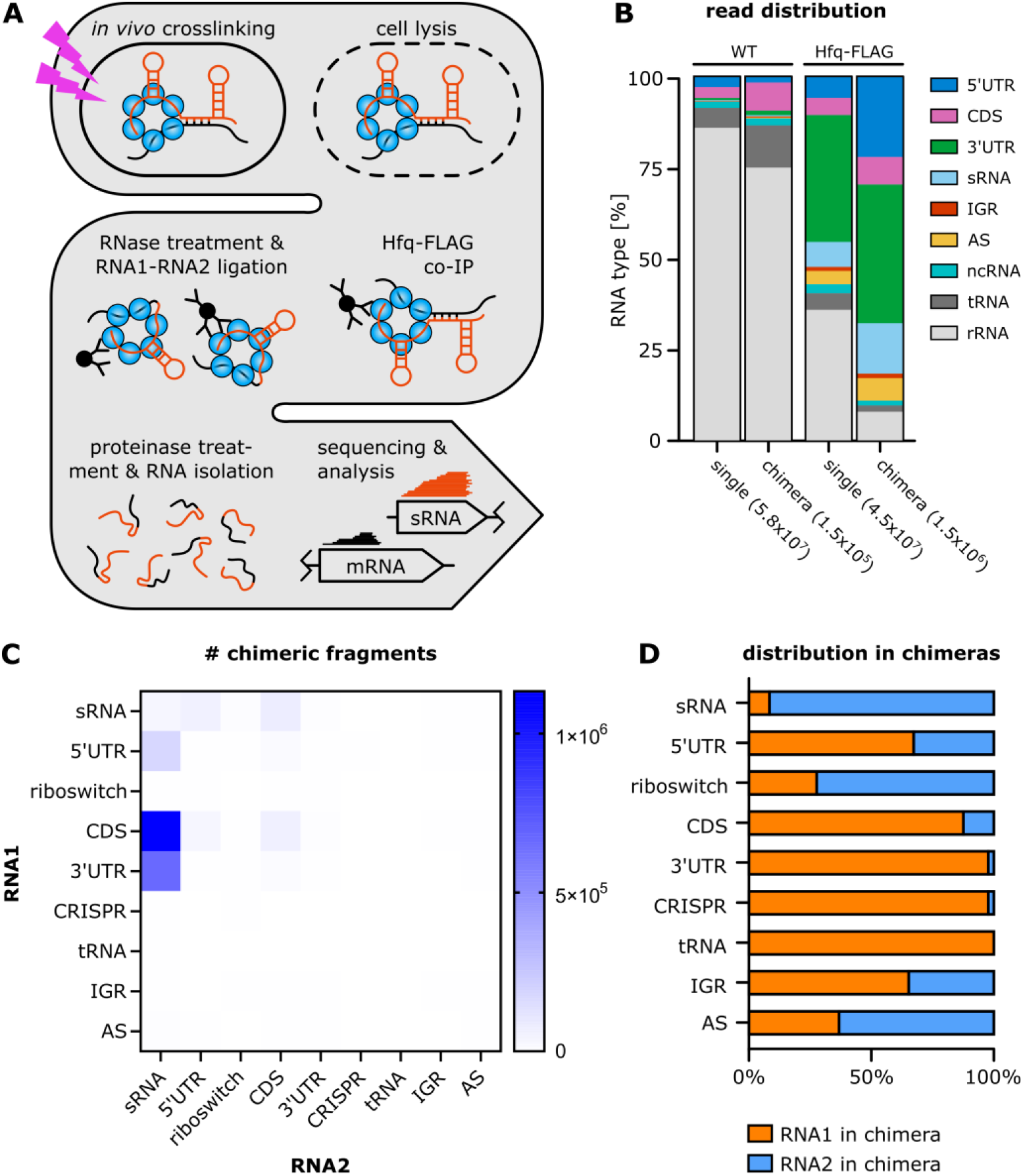
RIL-seq establishes Hfq as a platform for RNA-RNA interactions in *C. difficile*. **(A)** Schematic representation of the RIL-seq workflow. (B) Distribution of all reads, single and chimeric, across all RNA classes, comparing Hfq-FLAG and control (WT) strain (n=4 each). ncRNAs include riboswitches, tmRNA, SRP RNA, RNase P RNA and 6S RNA. All chimeras were included, without filtering for statistical significance or manual curation. (C)&(D) Distribution of RNA classes in chimeric fragments based on Supplementary Table 3, where RNA1 constitutes the 5’end and RNA2 the 3’end of a chimera. Only statistically relevant interactions (Fisher’s exact test ≤ 0.05) that are represented by at least 25 chimeric fragments are included.

### Hfq RIL-seq identifies novel sRNA candidates

Recent publications suggest that RIL-seq network data can be exploited to identify new sRNAs by taking into account unique features of sRNAs in general and sRNA RIL-seq chimeras in particular^31,36^. Accordingly, a high number of chimeric fragments mapping to a single RNA has been identified as a promising indicator of potential new sRNAs^36^. This is reflected in the formation of “interaction hubs” that consist of a dominating, single RNA interacting with a large number of unique RNAs^31^. By mapping all chimeric fragments to the *C. difficile* genome we could identify 24 interaction hubs formed by sRNA candidates that were previously unknown (n=15) or non-validated (n=9) sRNAs (Figure 2A, Supplementary Table 4) ^18,37,38^. Subsequent northern blot analysis confirmed the expression of six out of eight tested sRNA candidates (Figure 2B). Expression profiles of these sRNAs indicated expression mainly during late exponential/early stationary growth phase (Figure 2B), coinciding with the growth stage selected for our RIL-seq experiment. Accordingly, performing RIL-seq in distinct growth conditions has the potential to uncover novel sRNA candidates that have evaded previous detection approaches such as RIP-seq, due to its unique ability to reveal both Hfq-association and RNA-RNA interaction^24^.

**Figure 2:**
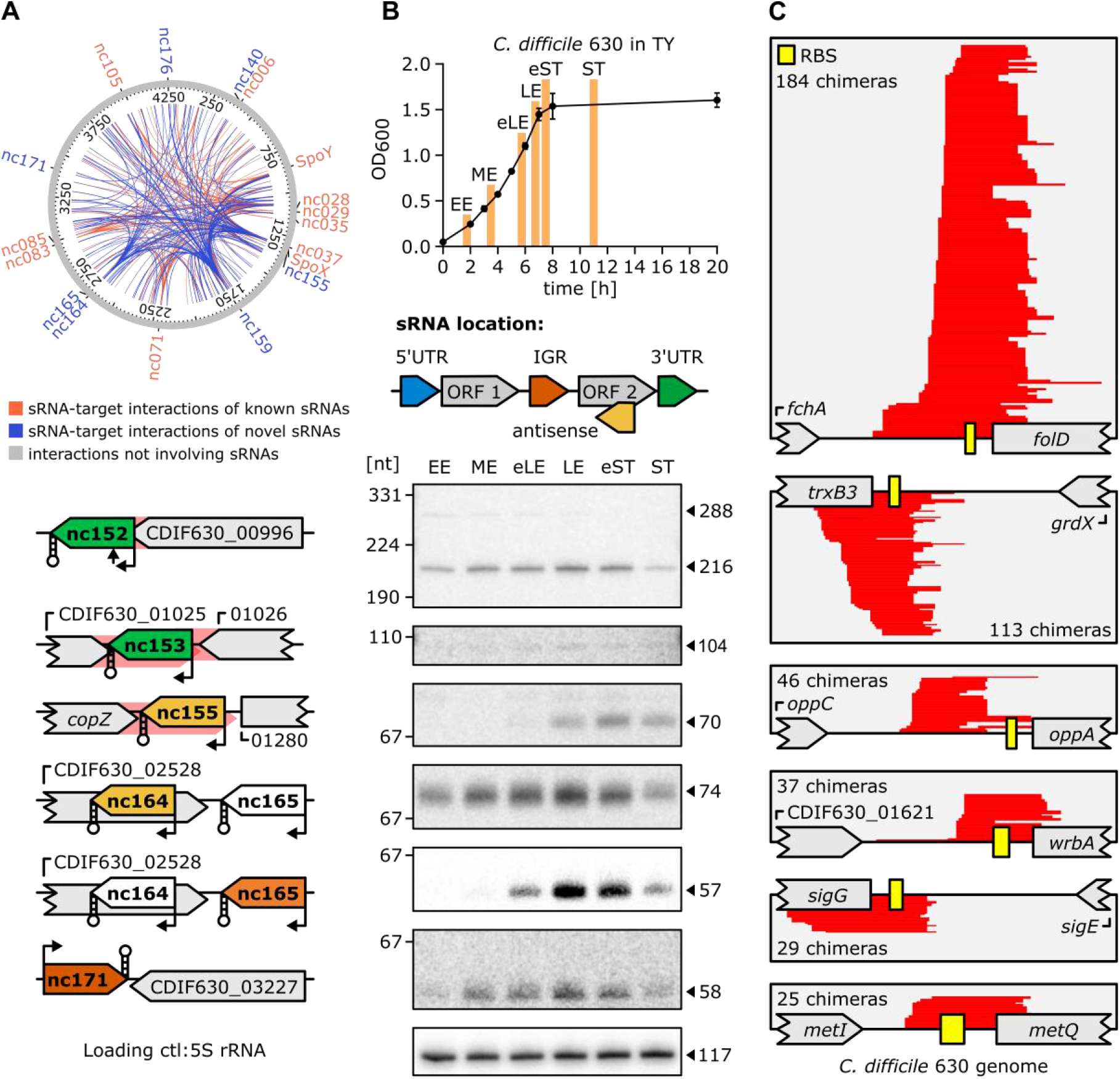
RIL-seq analysis facilitates annotation of novel sRNAs and reveals sRNA-mediated regulation of polycistronic transcripts. **(A)** Circos plot of all RIL-seq interactions that are represented by ≥ 200 chimeric fragments, mapped to the C. difficile 630 chromosome. Interaction hubs characterized by sRNAs with ≥ 20 unique interactions are labelled. Interactions involving known sRNAs are marked in orange, while interactions involving new sRNAs are highlighted in blue. All other interactions are grey. (B) Northern blot validation of new sRNAs. Samples were taken in early exponential (EE), mid-exponential (ME), early late exponential (eLE), late exponential (LE), early stationary (eST) and stationary (ST) phase of growth. sRNAs are color-coded according to their genomic location, stem loop structures indicate intrinsic terminators, arrows TSSs or processing sites and red shadings untranslated regions. A representative image of three independent experiments is shown. (C) Coverage plots of several sRNA-IGR chimeras, where the IGR is located within poly-cistronic operons. Chimeric reads are highlighted in red; the RBS is depicted as a yellow box. The number of chimeras covering each interaction is provided.

### RIL-seq data suggests sRNA-mediated discoordinate regulation of operons in C. difficile

In addition to the vast number of chimeras representing sRNA-mRNA pairs (n=1046), sRNA encompassing interactions also included chimeras consisting of sRNA-sRNA (n=39) and sRNA-IGR ligations (n=24, Supplementary Table 2). While sRNA-sRNA pairs have been discussed as a pool of potential sRNA sponges, sRNA-IGR chimeras have not been investigated previously^24,31,39^. A detailed analysis of those interactions revealed that in several cases (n=8, 33%) “IGRs” represented non-coding regions in polycistronic mRNAs Figure 2C). To further understand the impact of sRNA-mediated regulation on operon expression in *C. difficile*, we compared our RIL-seq data with previously published operon annotations^18^. We found that 383 RIL-seq chimeras mapped to 170 out of 400 known operons. Of these 383 chimeras, 98 constituted sRNAs interacting with intra-operon ribosome binding site (RBS) regions (25 nt up- and 20 nt downstream of the respective start codon), indicating potentially widespread coordinate and discoordinate regulation of polycistronic mRNAs in *C. difficile* (Supplementary Table 5). While sRNA-mediated coordinate regulation of entire operons is more common, there are fewer reports of sRNAs targeting individual genes in polycistronic mRNAs, thereby only affecting a subset of genes within an operon (discoordinate regulation) ^40,41^. For instance in *E. coli*, RyhB targets the *iscRSUA* operon, selectively inhibiting translation of *iscS* and resulting in the degradation of the *iscSUA* part^42,43^. Hence, discoordinate regulation of operons allows bacteria to selectively produce operon components, *e,g*. when only a specific gene product is needed in a given conditions^40^. In line, our RIL-seq dataset comprised chimeras formed by the RBS region of the sporulation-specific sigma factor *sigG* and the newly annotated sRNA CDIF630nc_161 (Figure 2C). *sigG* encodes the forespore-specific late sporulation sigma factor σ^G^ and constitutes the last gene in an operon formed by two additional sporulation specific genes, including *sigE* directly upstream of *sigG*^44^. In contrast to σ^G^, the *sigE* encoded sigma factor σ^E^ is active in the mother cell during early sporulation. Accordingly, both sigma factors not only operate during different stages of sporulation, but also in different compartments, and consequently require a tight regulation^44^. Hence, sRNA-mediated post-transcriptional regulation might finetune the sequential expression of both sigma factors to ensure correct spore development. Nevertheless, a more detailed analysis is needed to fully understand the nature and extent of these regulatory events in *C. difficile* not only on sporulation but on cellular processes in general.

### The master regulator of sporulation, Spo0A, is a central target of sRNA-based regulation

Interestingly, *sigG* was only one among several sporulation specific genes enriched in our RIL-seq dataset. Additional sporulation related genes included *sigE, sleB, spoIIAB, spoIVA* and *spo0A*, encoding the master regulator of sporulation (Supplementary Table 2)^10^. The latter was of particular interest since chimeras comprising *spo0A* and the sRNA nc020 as well as *spo0A* and sRNA nc038 were among the top 5 most abundant RIL-seq interactions in the entire dataset (>20,000 chimeras respectively). We decided to investigate these interactions in more detail and renamed both sRNAs to SpoY (nc020) and SpoX (nc038), to reflect their involvement in sporulation. SpoY is a 5’UTR derived sRNA, sharing its transcription start site with CDIF630_00827, which encodes a protein of unknown function (Figure 3A&B). Northern blot analysis indicated a complex expression profile with the highest SpoY expression during the early- and mid-exponential growth phases, decreasing levels towards early stationary growth and increasing expression following entry into stationary phase (Figure 3C). MEME analysis of SpoY RIL-seq chimeras, including the *spo0A* interaction, suggested that SpoY preferably binds mRNA 5’UTRs at a G-rich target motif that resembles the RBS (Figure 3A&B, Supplementary Figure 3A&B)^45^. Indeed, i*n silico* predictions of RNA-RNA interactions performed with IntaRNA and RNAcofold, indicated that SpoY binding blocks the *spo0A* start codon region, which typically leads to translational inhibition (Figure 3D, Supplementary Figure 4A&B)^46,47^.

**Figure 3:**
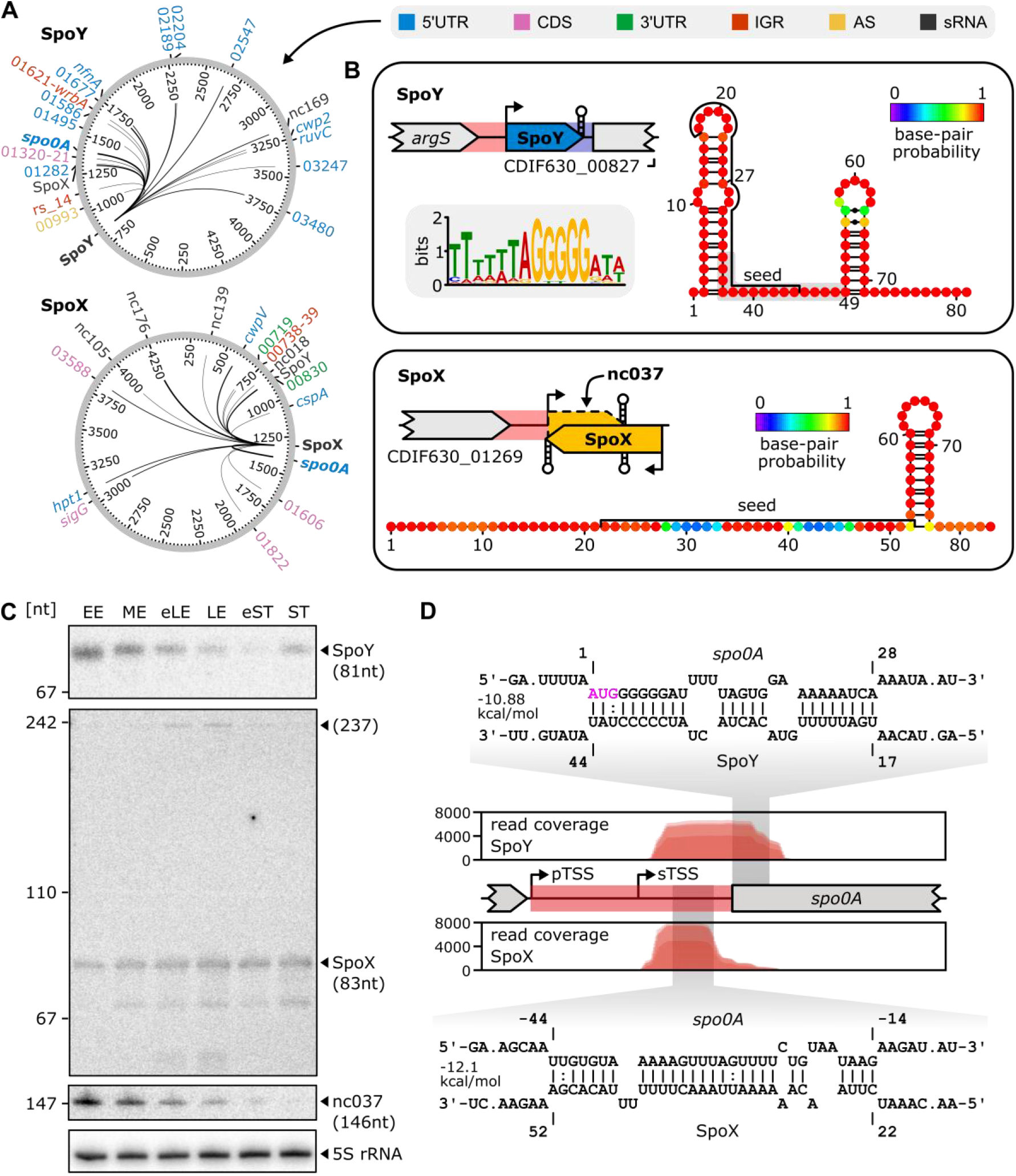
RIL-seq reveals *spo0A* as a target of sRNA-mediated post-transcriptional regulation. **(A)** Circos plots highlighting the SpoY and SpoX interactome, only interactions supported by ≥25 chimeras are included. Target types are discriminated by color. Edge strength correlates with the number of chimeras supporting an individual interaction. (B) Genomic location and predicted secondary structure (RNAfold^**80**^) of SpoY and SpoX are provided. Seed regions relevant for *spo0A* interaction were predicted *in silico* (IntaRNA^47^) and are labeled in the secondary structure. For both sRNAs, target sequences were extracted from the RIL-seq data and uploaded to MEME^45^, resulting in the successful identification of a common SpoY target motif (present in 24/28 target sequences). (C) Northern blot validation of SpoY, SpoX and nc037 expression in early exponential (EE), mid-exponential (ME), early late exponential (eLE), late exponential (LE), early stationary (eST) and stationary (ST) phase of growth in TY medium. 5S rRNA served as a loading ctl. A representative image of three independent experiments is shown. (D) Read coverage of spo0A by SpoY-spo0A (top) and SpoX-spo0A (bottom) chimeric reads. Base pairing information and location of predicted interaction sites (IntaRNA^47^) are highlighted. The spo0A nucleotide position is calculated relative to the spo0A start codon (highlighted in pink). The sigA dependent primary transcription start site (pTSS) and sigH dependent secondary TSS are marked.

In contrast to SpoY, SpoX is encoded partially antisense to another putative sRNA, nc037 (Figure 3B). Interestingly, previously published RNA-seq data revealed an intrinsic terminator within the SpoX sequence, resulting in a short (83 nt) and long isoform (237 nt, Figure 3B, Supplementary Figure 3B)^18^. The long isoform is encoded antisense to nc037, while the short isoform terminates prior to the overlapping region. According to northern blot analysis, the short SpoX isoform is more prevalent and uniformly expressed through all growth phases, while the long isoform is only present during the late exponential growth phase (Figure 3C). In contrast, expression of the putative sRNA nc037 is mostly anticorrelated (Figure 3C), potentially influencing expression of the long SpoX isoform. Further analyses will be necessary to assess the impact of nc037 on SpoX expression. The target spectrum of SpoX is diverse, including several CDSs and sRNAs in addition to the *spo0A* 5’UTR (Figure 3A, Supplementary Figure 3A). Consequently, identification of a conserved target motif using MEME was not successful^45^. According to IntaRNA analysis and the peak profile of SpoX-*spo0A* chimeric reads mapping to the *spo0A* 5’UTR, SpoX binds further upstream in the *spo0A* 5’UTR (Figure 3D)^47^. *In silico* predictions of the secondary structures of SpoX and the *spo0A* mRNA upon dimer formation suggested that SpoX-*spo0A* base pairing disrupts the *spo0A* 5’UTR secondary structure, potentially rendering the RBS more accessible to ribosome binding (Supplementary Figure 4A&B). Of note, the predicted seed region involved in *spo0A* interaction was located at the beginning of the SpoX sRNA and therefore present in both SpoX isoforms (Figure 3B&D, Supplementary Figure 3B)^47^. However, SpoX-*spo0A* chimeric reads solely mapped to the short isoform, which suggested that the short rather than the long version of SpoX predominantly binds *spo0A* (Supplementary Figure 3B). In summary, although the RIL-seq data revealed that both SpoX and SpoY interact with *spo0A*, their distinct interaction sites and chimeric read profiles suggested that the regulatory mechanisms applied by SpoX and SpoY differ.

### SpoY and SpoX directly bind the spo0A mRNA in vitro and in vivo

To confirm the *in silico* predicted sRNA-*spo0A* interactions, we performed electrophoretic mobility shift assays (EMSAs), combining either SpoY or SpoX with the full-length *spo0A* 5’UTR and start of CDS (Supplementary Figure 4A). Considering the location of the SpoX seed region we decided to use the short SpoX isoform for *in vitro* experiments. Complex formation of SpoY-*spo0A* required high concentrations of *spo0A*, but clearly improved upon addition of purified Hfq (Figure 4A, Supplementary Figure 5B). The complex of SpoX and *spo0A* formed more efficiently, resulting in an apparent K_D_ of 8.7 nM (Figure 4A, Supplementary Figure 5C). Mutating the respective sRNA seed regions (SpoY*/SpoX*) completely abolished the interaction in both cases, while introducing compensatory mutations in the respective *spo0A* target regions (*spo0A*\*^*C*^) restored the complex formation, albeit not to WT levels (Figure 4A, Supplementary Figure 5A-C). In line probing analysis further corroborated these results. As shown in Figure 4B, duplex formation of 5′-end-labeled SpoY and *spo0A* protected SpoY from cleavage at positions 30-41, partially confirming the predicted interaction site. Base-pairing was even more apparent for SpoX, where a clear concentration dependent effect could be observed, protecting SpoX from spontaneous cleavage at positions 22-39 & 47-52 upon duplex formation with *spo0A* (Figure 4B, Figure 3D, Supplementary Figure 4B). Based on these results we conclude that SpoY and SpoX interact with *spo0A in vitro via* direct base-pairing at distinct target sites in the *spo0A* mRNA. To further verify these interactions and their impact on *spo0A* translation *in vivo*, we designed a translational reporter system, consisting of mCherry fused C-terminally to *spo0A* (Supplementary Figure 5D). The *spo0A* fusion construct was either expressed alone (*p*[*spo0A*]) or in combination with one of the two sRNAs (*p*[SpoY/SpoX-*spo0A*]) in the respective sRNA deletion mutant. In contrast to the *in vitro* approaches described above, the long SpoX isoform was used for all reporter assays to fully reflect the *in vivo* situation, including potential regulation by nc037. Constitutive co-expression of SpoY and the mCherry fusion construct significantly decreased fluorescence as compared to the ΔSpoY-*p*[*spo0A*] control (ctl), demonstrating that SpoY inhibits *spo0A* translation (Figure 4C). Indeed, mutating the SpoY seed region (SpoY*) eliminated this inhibitory effect, while introducing the corresponding compensatory mutations in *spo0A* (*spo0A*\*^*C*^) restored the phenotype (Figure 4C, Supplementary Figure 4D). Interestingly, SpoX had the opposite effect on *spo0A* translation, as co-expression of SpoX and the mCherry fusion resulted in an increase of *spo0A* translation and consequently mCherry fluorescence (Figure 4C). *spo0A t*ranslation was restored to WT levels when co-expressing a SpoX* seed region mutant, however, introducing compensatory mutations in the *spo0A* 5’UTR did not restore the positive effect on translation. It is possible that the compensatory mutations interfere with the *spo0A* 5’UTR secondary structure, thereby preventing SpoX mediated opening of the *spo0A* 5’UTR to ribosome binding, as suggested above (Supplementary Figure 4B). Overall, we were able to confirm that SpoY and SpoX directly base-pair with the *spo0A* mRNA *in vivo*, resulting in translational repression of *spo0A* by SpoY and increased translation of *spo0A* upon interaction with SpoX.

**Figure 4:**
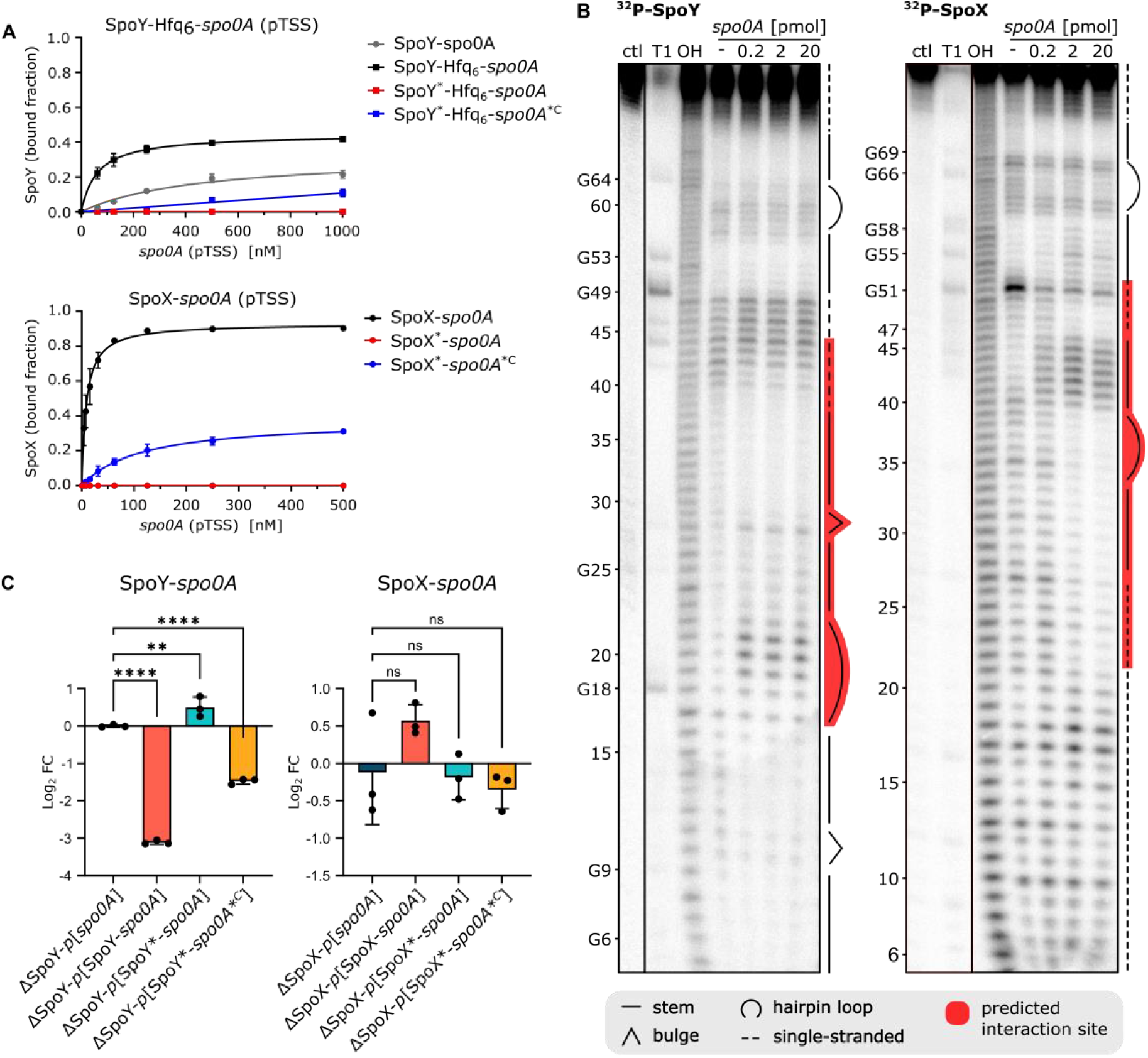
SpoY and SpoX directly interact with *spo0A in vitro* and *in vivo*. **(A)** Quantification of EMSAs (n=3, Supplementary Figure 5B&C) performed with either ^32^P-labeled SpoY or SpoX (short isoform) with increasing concentrations of the *spo0A* target region, respectively. Purified Hfq was added to facilitate SpoY-*spo0A* complex formation. Mutating the respective sRNA seed region (Supplementary Figure 5A, SpoY*/SpoX*) abolished the interaction, while introducing compensatory mutations into the *spo0A* target region (*spo0A\**^*C*^) slightly rescued the complex formation. **(B)** In-line probing of 0.2 pmol of ^32^P-labeled SpoY and SpoX in the absence (lane 4) or presence of increasing concentrations (lane 5-7) of the *spo0A* target region. RNase T1 and alkali-digested (OH) SpoY and SpoX serve as ladders respectively. Secondary structure and predicted seed region are highlighted. A representative image of three independent experiments is shown. **(C)** mCherry fluorescence of translational fusion constructs (n=3 replicates, Supplementary Figure 5D) expressed in the respective sRNA knock-out background. Fluorescence intensity was normalized to that of the respective *p*[*spo0A*] ctl. Mutating the sRNA seed regions (Supplementary Figure 5A, SpoY*/SpoX*) abolished the impact on *spo0A* translation, while introducing compensatory mutations into the *spo0A* target region (*spo0A\**^*C*^) partially rescued the effect. Ordinary one-way ANOVA with Dunnett’s multiple comparison test was used to calculate statistical significance. Not significant (ns), *P* ≥ 0.05; (**) *P* < 0.001 to 0.01; (****) *P* < 0.0001.

### Post-transcriptional regulation of spo0A by SpoY and SpoX has opposing effects on sporulation

Considering the evident alteration of *spo0A* translation, we hypothesized that SpoY and SpoX impact Spo0A protein levels *in vivo*. Accordingly, we performed western blot analysis to compare Spo0A protein levels in a WT strain (*p*[ctl]) to those in the SpoY and SpoX deletion mutants (e.g., ΔSpoY-*p*[ctl]), or to strains constitutively over-expressing the respective sRNA (e.g., ΔSpoY-*p*[SpoY]). Although there was no effect on Spo0A in a SpoY deletion mutant, overexpression of SpoY resulted in a significant decrease in Spo0A protein levels (∼2.5-fold, Figure 5A), confirming that SpoY inhibits *spo0A* translation. In contrast, deleting SpoX slightly decreased Spo0A levels, while over-expression of SpoX restored the Spo0A signal to WT levels. These results corroborated our model, in which SpoX positively impacts Spo0A translation by base-pairing to the *spo0A* 5’UTR. Although SpoX and SpoY regulate *spo0A* translation, changes in Spo0A levels might not directly translate into changes in Spo0A activity, as Spo0A requires additional activation *via* phosphorylation (Spo0A-P)^8^. To evaluate if SpoX and SpoY mediated changes in Spo0A levels correlated with Spo0A-P activity, we analyzed transcription of several sporulation-specific genes that operate downstream of Spo0A-P (Figure 6). Besides *spo0A*, transcript levels of *sigE, sigF, spoIV* (σ^E^ regulon), *spoIIQ* (σ^F^ regulon), *sigK, sigG* and *sspA* (σ^G^ regulon) were measured *via* qRT-PCR^44,48^. In accordance with our previous results, SpoY overexpression had a negative effect on transcript levels of all tested genes, whereas SpoY deletion had either no effect (*spo0A, sigE, sigF*) or resulted in an increase of transcript abundance. In contrast, deletion of SpoX reduced transcript levels of all tested genes, while SpoX overexpression partially restored transcript abundance to WT levels. Taken together, the observed changes in expression of sporulation-specific genes suggested that modulation of Spo0A levels by SpoY leads to an overall downregulation of the sporulation cascade, while activity of SpoX results in upregulation of spore formation.

**Figure 5:**
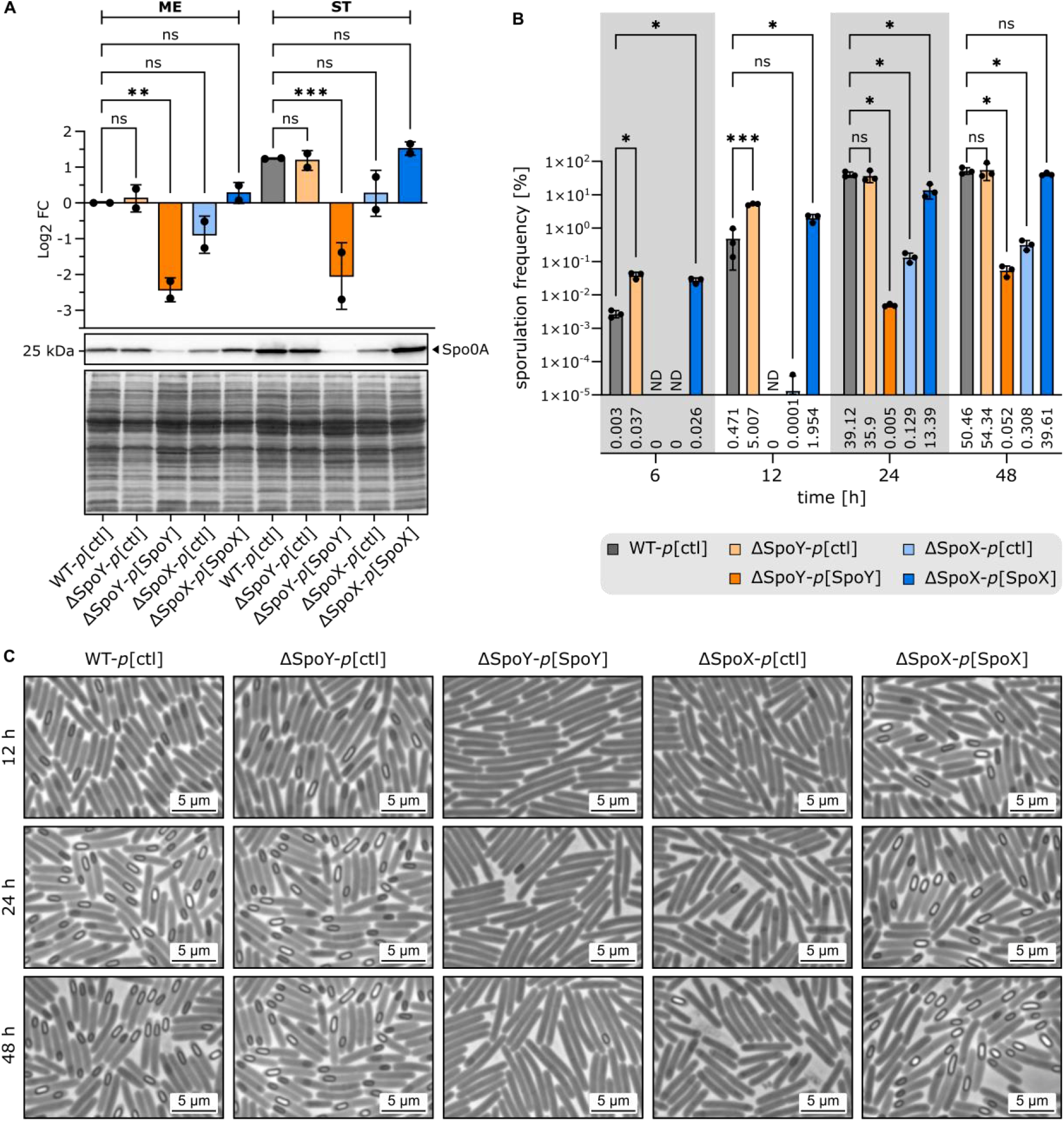
sRNA-mediated regulation of *spo0A* affects Spo0A levels and sporulation frequencies. **(A)** Western blot analysis comparing Spo0A protein levels in a WT strain (*p*[ctl]), sRNA knock-out mutants (ΔSpoY/ΔSpoX*-p*[ctl]) and strains constitutively expressing the respective sRNA (ΔSpoY/ΔSpoX*-p*[ΔSpoY/ΔSpoX]). Equal optical density (OD) units of total cell lysates (n=2 replicates) were loaded from strains gown till mid-exponential (ME) or stationary (ST) phase of growth. Band intensities were measured and Log_2_ fold changes were calculated relative to the WT (ME). Western blot membranes were incubated with anti-Spo0A antibody. Ponceau S staining of the blotting membrane served as loading control (bottom panel). **(B)** Sporulation frequencies of strains (n=3 replicates) described in (A) at 6 h, 12 h, 24 h and 48 h post inoculation of 70:30 liquid sporulation medium. ND: not determined – no viable spores. **(C)** Representative phase-contrast images of strains (n=3 replicates) described in (A) and treated as described in (B). Sporulation frequencies obtained from phase-contrast microscopy are provided in Supplementary Figure 6. To calculate statistical significance, an ordinary one-way ANOVA with Dunnett’s multiple comparison test was applied in (A) and a 2-way ANOVA with Dunnett’s multiple comparison test in (B). Not significant (ns), *P* ≥ 0.05; (*) *P* 0.01 to 0.05; (**) *P* < 0.001 to 0.01; (***) *P* < 0.0001 to 0.001.

**Figure 6:**
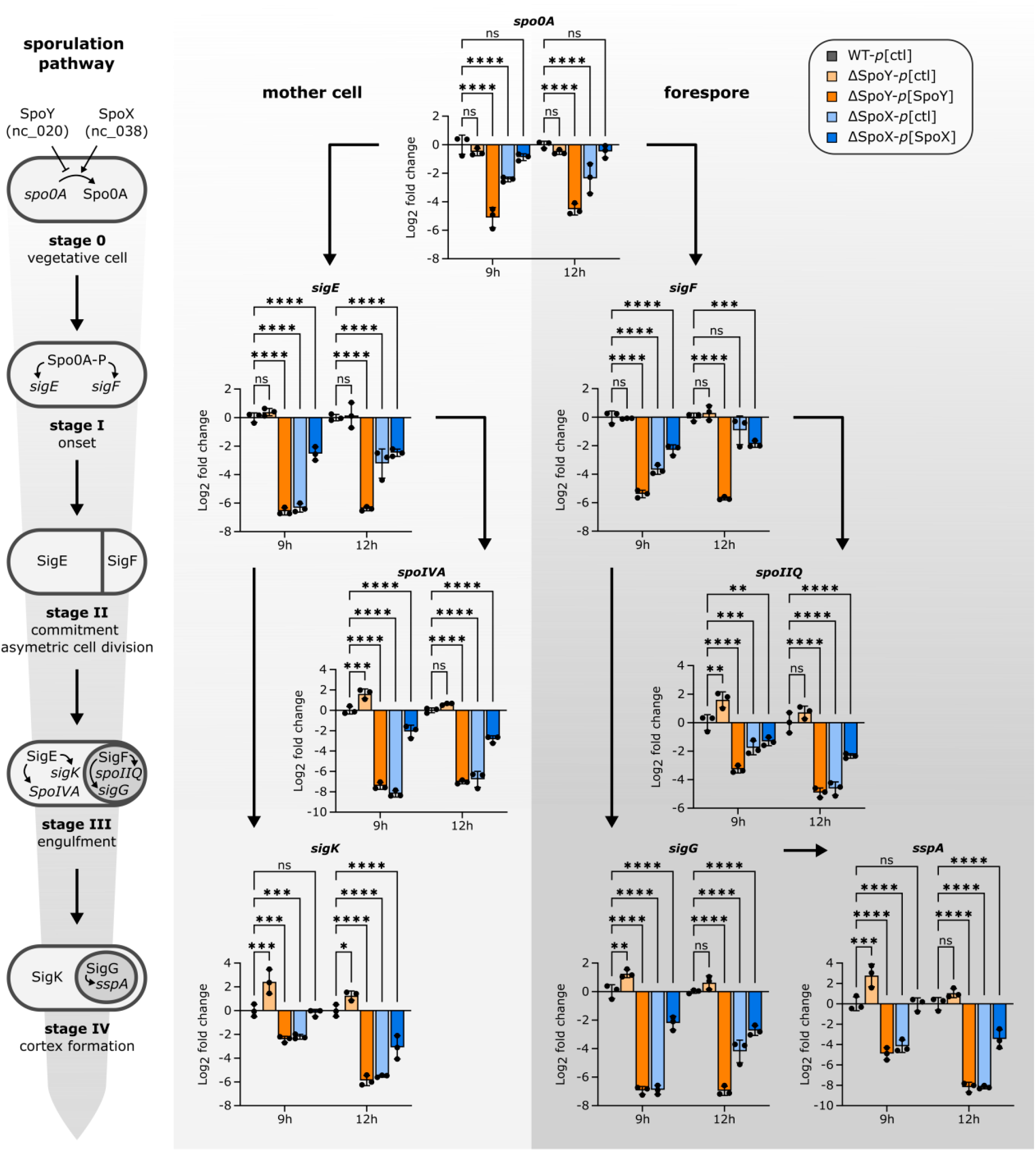
sRNA mediated post-transcriptional regulation of *spo0A* affects sporulation-specific gene expression. Relative transcript levels of genes encoding sporulation-specific sigma factors and their respective regulon in *C. difficile* 630 WT (*p*[ctl]), sRNA knock-out mutants (ΔSpoY/ΔSpoX-*p*[ctl]) and strains constitutively expressing the respective sRNA (ΔSpoY/ΔSpoX-*p*[ΔSpoY/ΔSpoX]). A schematic representation of the first four stages of sporulation in *C. difficile* is given on the left. RNA was extracted from samples (n = 3 replicates) taken at 9 h and 12 h post induction of sporulation on 70:30 sporulation plates. Log_2_ fold changes were calculated relative to the WT. Statistical significance was determined using two-way ANOVA with Dunnett’s multiple comparison test. Not significant (ns), *P* ≥ 0.05; (*) *P* 0.01 to 0.05; (**) *P* 0.001 to 0.01; (***) *P* 0.0001 to 0.001; (****) *P* < 0.0001.

To further assess the impact of SpoY and SpoX on sporulation, we determined sporulation frequencies of WT and mutant strains. As shown in Figure 5B, SpoY over-expression and SpoX deletion resulted in significantly reduced sporulation frequencies. SpoY deletion and SpoX over-expression on the other hand, slightly increased sporulation, particularly during early time points (6 h & 12 h). These observations were confirmed by phase contrast microscopy, revealing comparable phenotypes (Figure 5C, Supplementary Figure 6). Hence, our data show, that SpoY and SpoX not only affect *spo0A* translation in an inverse manner, but consequently influence gene expression of sporulation-specific genes and ultimately sporulation frequencies.

### SpoY and SpoX impact C. difficile gut colonization in a mouse model of C. difficile infection

Considering the marked impact of SpoY and SpoX deletion on sporulation in *C. difficile*, as well as their extended interactome, we decided to monitor the effect of SpoY and SpoX deletion in a mouse model of *C. difficile* infection (Figure 7A). Overall, a delayed onset of disease in mice challenged with the sRNA deletion strains was observed, as ΔSpoY and ΔSpoX infected mice showed a delayed body weight loss compared to mice treated with *C. difficile* WT (Figure 7B). Nevertheless, colon shortening on day 7 was equally severe in mice challenged with *C. difficile* WT, ΔSpoY or ΔSpoX suggesting similar levels of toxin production and consequently disease severity over the course of infection (Figure 7C). Initial colonization was comparable in ΔSpoY, ΔSpoX and WT treated mice, as no difference in vegetative cells or spores was observed at day 1 post infection (Figure 7D). However, lower CFUs of spores and vegetative cells were recovered from feces of ΔSpoY and ΔSpoX infected mice at days 3, 5 and 7 post infection, compared to mice challenged with *C. difficile* WT. Accordingly, mice infected with ΔSpoY or ΔSpoX strains exhibited an accelerated *C. difficile* clearance rate (Figure 7E). This was particularly evident in ΔSpoY treated mice starting from day 3, while the bacterial burden in mice infected with ΔSpoX only decreased at day 5 (vegetative cells) and day 7 (vegetative cells and spores) compared to WT treated mice. Generally, the lower spore counts in ΔSpoY or ΔSpoX infected mice were paralleled by lower vegetative cell counts, revealing a global effect of SpoY and SpoX deletion on *C. difficile* gut colonization rather than on sporulation alone (Figure 7D). Taken together, the impact of SpoY and SpoX on intestinal pathogenesis suggested that their regulatory functions extend beyond regulating Spo0A protein levels and likely include additional regulatory targets that contribute to intestinal colonization.

**Figure 7:**
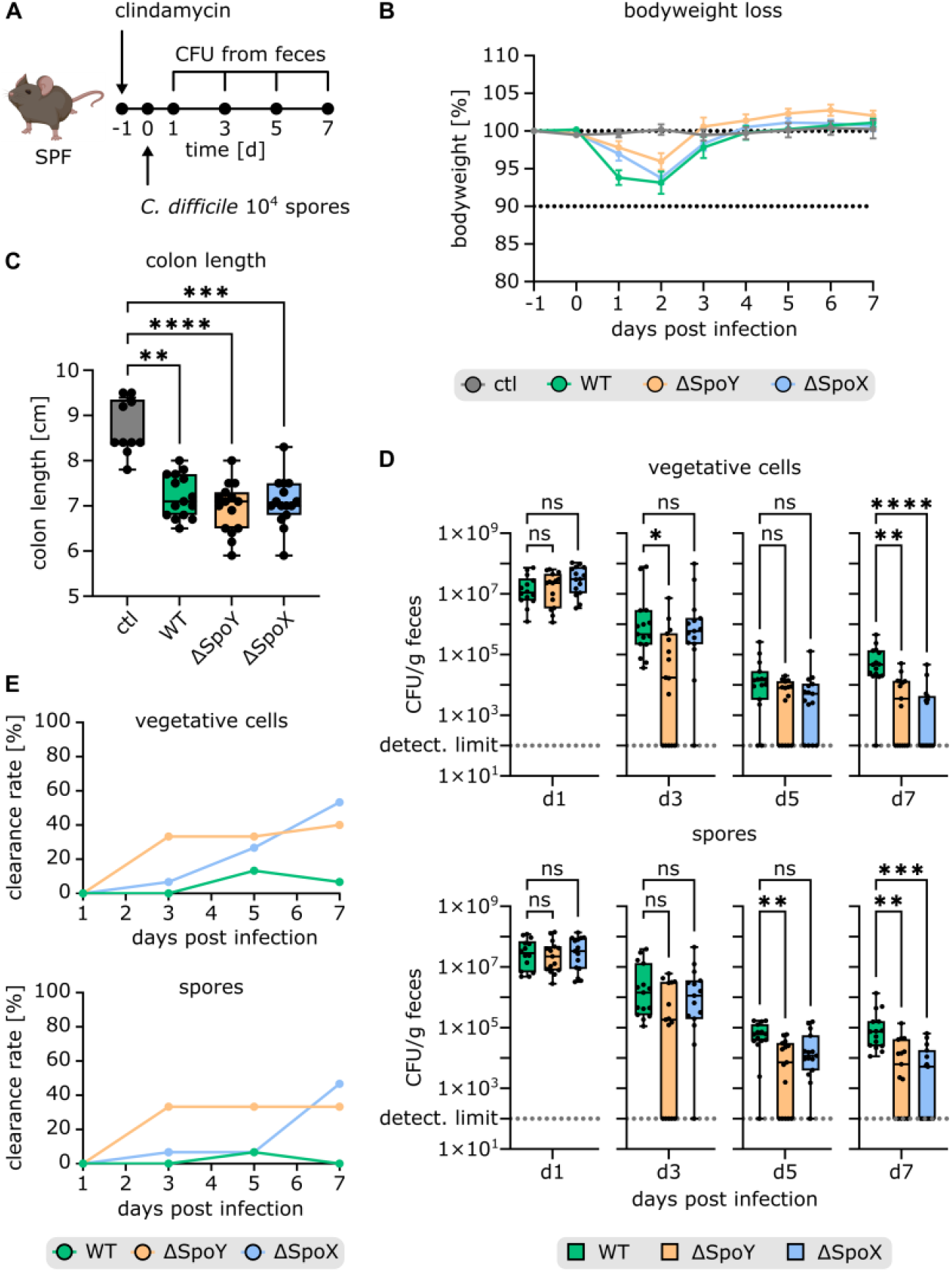
SpoY and SpoX deletion affects *C. difficile* gut colonization and spore burden in a mouse model of *C. difficile* infection. **(A)** Schematic representation of the mouse model of *C. difficile* infection (partially created with BioRender.com). SPF mice were treated with clindamycin 24 h prior to infection, administered via intraperitoneal injection to induce susceptibility to *C. difficile* infection^79^. Following antibiotic treatment, groups of mice were infected with 10^4^ spores of *C. difficile* 630 WT (n=15), ΔSpoY (n=15) or ΔSpoX (n=15) via oral gavage. Mice were monitored for disease between days 0 and 7 post infection. Fecal samples were collected at indicated time points to determine pathogen burden. **(B)** Body weight loss over the course of 7 days (data points and error bars represent mean ± standard error of the mean) as well as **(C)** final colon length at day 7 post infection are indicated. **(D)** Comparison of CFUs of *C. difficile* vegetative cells and spores in fecal pellets at different time points during infection. Replicates with CFUs below the detection limit were set to 100. **(E)** Clearance rate (number of mice below detection limit divided by the total number of mice) of SPF mice infected with *C. difficile* WT, ΔSpoY or ΔSpoX strains. Kruskal-Wallis was performed with Dunnett’s multiple comparison test to calculate statistical significance in (C)&(B). Not significant (ns), P ≥ 0.05; (*) P 0.01 to 0.05; (**) P < 0.001 to 0.01; (***) *P* 0.0001 to 0.001; (****) *P* < 0.0001.

## DISCUSSION

A plethora of research in human pathogens such as *Pseudomonas aeruginosa, S. enterica*, and *Vibrio cholerae* has highlighted the importance of sRNAs in regulating virulence pathways^49^. To uncover sRNA-mediated regulatory mechanisms that might shape *C. difficile* virulence, we performed RIL-seq during the onset of sporulation in *C. difficile*^28^. Endospore formation has been extensively studied, particularly in the model organism *B. subtilis* and is a tightly regulated process, defined by several sequential morphological stages (Figure 6)^50^. Although the sporulation cascade is generally conserved between *C. difficile* and other endospore-forming Firmicutes, there are some striking differences, most notably regarding sporulation initiation^13^. In *B. subtilis*, environmental signals that induce sporulation are channeled through a complex phosphorelay system consisting of several sensor kinases and phosphotransferases, culminating in the activation of Spo0A^51^. Phosphorylated Spo0A-P then initiates the sporulation process by activating the transcription of several key sporulation-specific genes^51^. Unlike *B. subtilis, C. difficile* does not encode an apparent intermediate phosphorelay system^8^. Although three putative sensor histidine kinases (PtpA-C) have been described to directly influence Spo0A phosphorylation, the overall process of Spo0A activation remains barely understood and points towards additional unknown mechanisms regulating Spo0A activity in *C. difficile*^8,52,53^.

In this study, we identified sRNA-mediated post-transcriptional regulation of *spo0A* translation as a new mechanism contributing to sporulation initiation in *C. difficile*. There are a few examples of sRNA-mediated regulation of sporulation in endospore forming Firmicutes. In *B. subtilis* sRNA SR1 inhibits translation of the histidine kinases *kinA* that transmits environmental signals, eventually resulting in phosphorylation of Spo0A^54^. Another example is the *virX* sRNA in *C. perfringens* that negatively regulates sporulation by repressing transcription of the early forespore-specific σ factor σ^F 55^. Furthermore, sRNA RCd1 in *C. difficile* inhibits production of the late mother cell-specific σ factor σ^K^ by preventing the excision of the prophage-like element that interrupts the *sigK* gene^17^. Here, we characterized two novel sRNAs, SpoY and SpoX, that function by directly binding and regulating the *spo0A* mRNA (Figure 4, Figure 3D). We could show that SpoY operates by base-pairing with the *spo0A* start codon region, thereby inhibiting *spo0A* translation, resulting in reduced sporulation frequencies when overexpressed (Figure 5B&C). In contrast, SpoX interaction with the *spo0A* 5’UTR results in an upregulation of *spo0A* translation and consequently sporulation, most likely through a change of the *spo0A* 5’UTR secondary structure upon base-pairing with SpoX. Analogous mechanisms have been described in the literature; a well-known example is the positive regulation of *rpoS*, the key regulator of general stress responses in *E. coli*^56^. In this case, several sRNAs (DsrA, RprA, and ArcZ) base-pair with the *rpoS* 5′ leader to expose the *rpoS* translational start site that is otherwise blocked by an inhibitory stem-loop structure^56^.

Of note, chimeras formed by *spo0A* and SpoX or SpoY are not the only *spo0A* interactions present in the dataset, albeit the most enriched. In fact, we found nine additional chimeras consisting of the *spo0A* mRNA ligated to an sRNA (n=8) or another mRNA (n=1, Supplementary Figure 7). Of these nine interactions, eight displayed a chimeric read profile comparable to SpoY, suggesting a similar mode of action. Accordingly, these sRNAs might substitute for the SpoY function, potentially explaining the minor sporulation phenotype of a SpoY deletion strain. Interestingly, *in silico* predicted interaction sites overlap with the RIL-seq peak profile for five of the detected *spo0A*-sRNA interactions, further corroborating the RIL-seq results (Figure 5, Supplementary Figure 7)^47^. There are known examples of mRNAs that are directly targeted by multiple sRNAs, most of which encode key regulators. For instance, *flhDS*, the master regulator of flagellar genes in *E. coli* interacts with five sRNAs (ArcZ, OmrA, OmrB, OxyS, and McaS) that base-pair with the *flhDS* 5’UTR resulting in either negative or positive regulation of motility ^57^. Similarly, biofilm formation is a central target of sRNA mediated regulation through base-pairing of seven sRNAs (OmrA/B, McaS, RprA, RydC, GcvB, and RybB) with the *csgD* 5’UTR, encoding a central regulator of curli formation in *E*. coli^58^. In line with these examples, *C. difficile* Spo0A might constitute another key regulator that is a central target of sRNA-mediated regulation. Accordingly, the various sRNAs interacting with *spo0A* may serve to integrate different environmental signals to modulate Spo0A expression, thus partially replacing the missing phosphorelay system that translates environmental cues into Spo0A activity in *B. subtilis*^8,51^. This hypothesis is further corroborated by the distinct expression profiles of SpoY and SpoX, which points to *spo0A* regulation during different growth phases (Figure 3C). SpoY most likely suppresses *spo0A* translation during early growth stages, in conditions that favor active growth rather than sporulation. In contrast, SpoX expression indicates that it exerts its positive effect on *spo0A* translation mostly in late exponential and early stationary phase when the sporulation process initiates. Accordingly, finding the specific growth conditions that lead to altered expression of sRNAs predicted to regulate *spo0A* will be vital to fully understand the nuanced post-transcriptional regulation of *spo0A*. Of note, Spo0A has been implicated in pathways other than sporulation, as Spo0A inactivation also results in decreased toxin production and biofilm formation in *C. difficile*^59,60^. Hence, sRNA-mediated regulation of *spo0A* might impact additional processes besides sporulation that could not be covered in this work and will require further investigation^59–61^.

Interestingly, some of the sRNAs interacting with *spo0A*, also formed chimeras with SpoX, including SpoY, nc105 and nc176 (Supplementary Figure 3A). It is possible that SpoX not only upregulates *spo0A* translation, but also sponges sRNAs that would otherwise inhibit *spo0A* translation. This could explain why the deletion of SpoX has such a pronounced effect on sporulation frequencies (Figure 5). Alternatively, it is equally likely that SpoY, nc105 and nc176 sRNA regulate SpoX activity, preventing positive regulation of sporulation in conditions favoring active proliferation. sRNA sponges generally act by sequestering a target sRNA, thereby preventing the sRNA-target interaction, an effect that depends on the stoichiometry between the sponge, sRNA, and mRNA^62^. Reports of sRNAs simultaneously acting as sRNA sponge and mRNA regulator have been published previously, supporting this hypothesis^62^. For example ArcZ and CyaR, both known regulators of the *rpoS* mRNA, also interact with each other, as ArcZ overexpression reduces CyaR steady state levels and upregulates CyaR targets^63^. Further research is necessary to verify these SpoX-sRNA interactions and decipher if, when and how these interactions impact Spo0A activity^62^.

Although the interactions of SpoY and SpoX with *spo0A* and with each other have been discussed above, it is important to consider that both sRNAs also interact with additional targets (Supplementary Figure 3A). For example, SpoY also targets *cwp2*, a cell wall protein known to affect cellular adherence *in vitro*^64^. Furthermore, *cwpV* was found among the mRNAs targeted by SpoX, and encodes a cell wall protein described to promote cell aggregation^65^. Consequently, deleting or overexpressing either sRNA *in vivo* most likely affects processes besides sporulation and might explain the global effect of both sRNA deletion strains in the mouse model of *C. difficile* infection (Figure 7). In both cases, sRNA deletion resulted in a reduction of spore and vegetative cell shedding compared to mice infected with *C. difficile* WT, pointing to a broader effect on disease development.

In summary, the application of Hfq RIL-seq to *C. difficile* has revealed a global view of extensive Hfq-mediated RNA interactions (Figure 1B, Supplementary Table 2)^30^. Although we have barely scratched the surface of sRNA mediated regulation in *C. difficile*, our RIL-seq data represents a starting point for the characterization of additional processes modulated by sRNAs. In this work, we uncovered a new layer of post-transcriptional regulation in *C. difficile* hinting at a complex sRNA network regulating sporulation in this important human pathogen. Given the low conservation of mechanisms governing sporulation initiation these results might open an interesting avenue for potential therapeutic targets to counteract CDI. In fact, the use of antisense nucleic acids to selectively target species in a microbial community has gained attention as a promising alternative to conventional antibiotics^66,67^. Accordingly, mimicking or blocking the activity of sRNAs using antisense nucleic acid derivatives might represent an interesting alternative, especially considering the contribution of antibiotics to CDI recurrence^7,49^.

## METHODS

### Bacterial strains and growth conditions

A complete list of all *C. difficil*e and *E. coli* strains used in this study is provided in Supplementary Table 6. *C. difficile* 630 cultures were routinely grown anaerobically inside a Coy chamber (85% N_2_, 10% H_2_ and 5% CO_2_) in Brain Heart Infusion (BHI) broth or on BHI agar plates (1.5% agar) unless stated otherwise. If necessary, antibiotics were added to the medium at the following concentrations: thiamphenicol (TAP) 15 μg/ml, cycloserine (CS) 250 μg/ml, cefoxitin (FOX) 8 μg/ml. *E. coli* cultures were propagated aerobically in Luria-Bertani (LB) broth (10 g/l tryptone, 5 g/l yeast extract, 10 g/l NaCl) or on LB agar plates (1.5% agar) supplemented with chloramphenicol (CHL, 20 μg/ml). *E. coli* strain Top10 (Invitrogen) served as a recipient for all cloning procedures, and *E. coli* CA434 (HB101 carrying the IncPβ conjugative plasmid R702) was used as donor strain for plasmid conjugations into *C. difficile* 630.

### Plasmid construction

All plasmids and DNA oligonucleotides used in this study are listed in Supplementary Table 6 and 8, respectively. *E. coli* TOP10 was used for plasmid propagation according to standard procedures^68^.

#### pFF-53 - plasmid for generating a *hfq*::3×FLAG strain

For insertion of a C-terminal *hfq::*3×FLAG-tag, allelic exchange cassettes were designed with approximately 1.2 kb of homology to the chromosomal sequence flanking the up- and downstream regions of the *hfq* stop codon. Both homology regions were PCR-amplified from *C. difficile* 630 using high fidelity Fusion Polymerase (Mobidiag) with 5% DMSO, FFO-364/-365 and FFO-368/-369. The resulting fragments were gel purified with NucleoSpin Gel and PCR Clean-Up Kit (Macherey-Nagel). The 3×FLAG-tag (DYKDHDGDYKDHDIDYKDDDDK) was similarly amplified and purified with FFO-366/-367, using the previously published pFF-12 as a template.^18^ Insert assembly and ligation into PCR-linearized pJAK184 (FFO-362/-363) was achieved via Gibson Assembly (Gibson Assembly ® Master Mix, New England BioLabs) according to the manufacturer’s instructions, resulting in pFF-53^18^.

#### pFF-162/-163/-164/-245/-166/-248/-247 – plasmids for *in vitro* transcriptions

SpoY (FFO-958/-959), SpoY* (FFO-958/-960), SpoX (short isoform, FFO-961/-962), SpoX* (short isoform, FFO-1261/-1262) and *spo0A* (5’UTR starting from pTSS and first 84 nt of coding region, FFO-964/-966) were PCR-amplified from *C. difficile* 630 using Fusion Polymerase (Mobidiag), adding a 5′ overhang comprising the T7-promoter sequence (5′-GTTTTTTTTAATACGACTCACTATAGGG). For inserting SpoY* compensatory mutations in *spo0A, spo0A*\*^*C*^ was amplified in two parts using FFO-964/-1268 and FFO-1269-966, before joining both fragments *via* SOEing PCR with FFO-964/-966. SpoX* compensatory mutations were inserted similarly using FFO-964/1259 and FFO-1267/-966, followed by SOEing PCR with FFO-964/-966. In each case, PCR products were gel purified as described above. Subsequently, 3’-adenine overhangs were added to all PCR product using Taq Polymerase (Biozym). The resulting fragments were cloned into the StrataClone TA-cloning vector and transformed into StrataClone SoloPack competent cells according to the manufacturer’s protocol (StrataClone PCR Cloning Kit, Agilent), resulting in pFF-162 (SpoY), pFF-163 (SpoY*), pFF-164 (SpoX), pFF-245 (SpoX*), pFF-166 (*spo0A*), pFF-248 (*spo0A*\*^*C*^ SpoY*) and pFF-247 (*spo0A*\*^*C*^ SpoX*).

#### pFF-170/-171 – plasmids for generating SpoY and SpoX deletion mutants

For deletion of SpoY and SpoX, allelic exchange cassettes were designed with approximately 1.2 kb of homology to the chromosomal sequence flanking the deletion sites of SpoY and SpoX. To avoid polar effects on genes or sRNAs encoded adjacent to SpoY or antisense to SpoX (Figure 3B), the deleted region was restricted to nucleotide 11-46 in case of SpoY, and nucleotide 1-83 of SpoX in addition to 40 nt upstream of SpoX encompassing the SpoX promoter region. Homology arms were PCR amplified, and gel purified as described above, using FFO-977/-978 and FFO-979/-980 for the SpoY homology arms, and FFO-985/-986 and FFO-987/-988 for amplification of the SpoX homology regions, respectively. The homology arms were joined *via* SOEing PCR resulting in one large fragment encompassing both homology regions, and a BamHI/SacI restriction site at the 5’/3’end for both, SpoY (FFO-977/-980) and SpoX (FFO-985/-988). Following restriction digest using BamHI and SacI, the fragments were mixed in a 3:1 ratio with an equally digested and gel purified pJAK112 and ligated overnight at 4 °C using T4 DNA ligase (Thermo Scientific), resulting in pFF-170 (SpoY deletion) and pFF-171 (SpoX deletion)^18^.

#### pFF-185/-186/-191/-254/-285/-187/-192/-260/-289/-207-translational fusion reporter

To discern the impact of sRNA-target interactions on target translation, we designed a translational fusion system based on the previously published pDSW1728 mCherryOpt plasmid, that was initially designed to study gene expression^69^. The full reporter system (RS) constitutively co-expresses an sRNA controlled by an *fdxA* promoter and the target 5’UTR fused to mCherryOpt, controlled by a *cwp2* promoter. Regulation of target translation *via* sRNAs is measured by comparing fluorescence of the strain expressing the full RS (*p*[sRNA-*target*]) to a control strain only expressing the sRNA (*p*[sRNA]) or the target 5’UTR fused to mCherryOpt (*p*[*target*]). All plasmids were designed to allow easy exchange of individual components *via* restriction digestion and are illustrated in Supplementary Figure 5D. sRNA expression was verified *via* northern blot analysis (Supplementary Figure E&F). For generating a *p*[*target*] plasmid, the *cwp2* promoter and the *spo0A* 5’UTR (including 75 nt of CDS) were PCR amplified and gel purified as described above from *C. difficile* 630. FFO-1004/-1000 and FFO-1001/-1002 were used, respectively, thereby adding NheI/XhoI restriction at the 5’/3’end of the *cwp2* promoter, and a XhoI/SacI restriction site at the 5’/3’end of the *spo0A* 5’UTR, respectively. mCherryOpt was PCR amplified from pDSW1728 with FFO-1056/-1057 starting with the second codon of the mCherryOpt CDS, adding a SacI restriction site directly upstream and preserving the BamHI restriction site at the 3’end. All components were subjected to restriction digest using the appropriate restriction enzymes and ligated into the NheI/BamHI digested pDSW1728 vector as described above, resulting in pFF-185 (*p*[spo0A]). To generate a *p*[sRNA] plasmid, the *fdxA* promoter and SpoY were PCR amplified from *C. difficile* 630 with FFO-995/-1005 and FFO-1006/-1007, respectively, inserting an NheI restriction site at the *fdxA* 5’end and a SpoY overlapping region at the 3’end. Inserting a restriction site upstream of SpoY was avoided to preserve the sRNA primary and secondary structure. Accordingly, an *fdxA* overlapping region was added to the SpoY 5’end and an XbaI restriction site at the 3’end followed by the *slpA* terminator and a BamHI restriction site to prevent readthrough. Both fragments were joined *via* SOEing PCR with FFO-995/-1007, NheI/BamHI digested and ligated into the NheI/BamHI digested pDSW1728 vector, resulting in pFF-186 (*p*[SpoY]). Finally, the *p*[sRNA-*target*] plasmid was generated by PCR amplifying the *cwp2*-*spo0A* 5’UTR-mCherryOpt construct from pFF-185, using FFO-999 and FFO-1057, thereby exchanging the 5’ NheI with an XbaI restriction site followed by the *slpA* terminator. The resulting fragment was digested with XbaI/BamHI and ligated into the equally digested pFF-186 yielding pFF-191 (*p*[SpoY-*spo0A*]). All remaining plasmids were generated by exchanging either sRNA or target of the plasmids described above. For SpoX constructs, *fdxA* and SpoX were PCR amplified using FFO-995/-1008 and FFO-1009/-1010. Both fragments were joined *via* SOEing PCR with FFO-995/-1010, digested with NheI/XbaI and ligated into NheI/XbaI digested pFF-186 and pFF-191, resulting in pFF-187 (*p*[SpoX]) and pFF-192 (*p*[SpoX-*spo0A*]). For SpoY* constructs, SpoY* was PCR amplified from pFF-248 with FFO-1006/-1007 and joined *via* SOEing PCR with the previously amplified *fdxA* promoter, using FFO-995/-1007. The SOEing product was NheI/XbaI digested and ligated into the equally digested pFF-191, yielding pFF-254 (*p*[SpoY*-*spo0A*]). Next *spo0A* harboring compensatory mutations was PCR amplified from pFF-248, using FFO-1001/-1002, digested with XhoI/SacI, and ligated into XhoI/SacI digested pFF-254 resulting in pFF-285 (*p*[SpoY*-*spo0A*\*^*C*^]). Generation of SpoX* constructs was achieved by first amplifying SpoX* in two fragments, inserting the seed region mutations with FFO-1264/-1262 and FFO-1263/-1010. Both fragments were then joined *via* SOEing PCR with FFO-1264/-1010, followed by a second SOEing PCR, combining the previously amplified *fdxA* promoter and the full length SpoX*, using FFO-955/-1010. The resulting product was NheI/XbaI digested and ligated into a similarly digested pFF-191, yielding pFF-260 (*p*[SpoX*-*spo0A*]). Finally, *spo0A* harboring compensatory mutations was PCR amplified from pFF-241, using FFO-1001/-1002, digested with XhoI/SacI, and ligated into XhoI/SacI digested pFF-260, resulting in pFF-289 (*p*[SpoX*-*spo0A*\*^*C*^]). In addition to the reporter system constructs, an empty control vector was generated by linearizing pFF-191 *via* PCR with FFO-994/-1205, adding an additional NheI restriction site at the 5’end. The resulting product was NheI digested and re-ligated yielding pFF-207 (*p*[ctl]).

### Plasmid conjugation

For conjugation purposes, plasmids were transformed using 80 μl of electro competent *E. coli* CA434 mixed with 100 – 500 ng of plasmids in a pre-chilled electroporation cuvette. Following electroporation with 1.8 kV, 200 Ω, 4-5 sec, cells were recovered for 4 h at 37 °C in 1 ml LB. Colonies harbouring the plasmid were selected on LB supplemented with CHL and confirmed *via* colony PCR. Conjugation was performed according to Kirk and Fagan (2016), as published previously^18,70^.

### Generation of deletion and insertion strains

Gene deletions were constructed using homologous recombination as previously published^18,71^. In short, plasmids were conjugated in *C. difficile* 630 strains as described above. Following conjugation, colonies were screened for the first recombination event *via* PCR. Positive recombinants were streaked on non-selective BHI, followed by incubation for 2 – 3 days. Growth was harvested using 900 μl 1xPBS, and 50 μl of a 10^−4^ and 10^−5^ dilution of the mixtures were streaked either on CDMM supplemented with 50 μg/ml fluorocytosine for pJAK112 derived plasmids, or on TY containing 4% w/v xylose for pJAK184 derived plasmids. Once colonies appeared, 8 – 15 were re-streaked to purity and tested for secondary recombination events *via* PCR. To test for plasmid loss, colonies were simultaneously streaked on selective plates containing TAP. Sanger sequencing was applied to confirm successful deletions and insertions.

### RIL-seq (RNA interaction by ligation and sequencing)

#### RIL-seq experimental procedure

RIL-seq was performed, following the original protocol published by Melamed and collogues with minor alterations^24,25^. Briefly, *C. difficile* 630 WT and Hfq-FLAG were grown in sterile filtered TY in four biological replicates until OD_600_ of ∼1.2^28,29^. Of each replicate, 100 ODs were harvested (4,500 x g, 15 min at 4 °C), resuspended in 10 ml ice-cold 1xPBS and irradiated with UV light (254 nm, 80,000 mJ/cm2). Following centrifugation (4,500 x g, 15 min at 4 °C), pellets were resuspended in 800 μl of ice-cold wash buffer (50 mM NaH_2_PO_4_, 300 mM NaCl, 0.05% Tween, 1:200 diluted protease inhibitor cocktail set III (Calbiochem)) supplemented with 0.1 U/μL of recombinant RNase inhibitor (wash buffer-RIn, Takara). Mechanical cell lysis was achieved by mixing each sample with 0.1 mm glass beads and grinding in a Retsch mixer mill at a frequency of 30/s for 5 min. The grinding was repeated four times; after each step the adaptors were place on ice for 2 min. The resulting supernatant was transferred to a new tube (17,000 x g, 2 min at 4 °C), 400 μl wash buffer-RIn was added to the glass beads and the grinding was repeated once more. For Hfq co-immunoprecipitations, the accumulated lysates were incubated with protein A/G magnetic beads (Thermo-Fisher) pre-coupled with anti-Flag antibody (M2 monoclonal antibody, Sigma-Aldrich) for 2 h at 4°C with rotation, followed by three washing steps with wash buffer-RIn. Next, samples were treated with 480 μl of wash buffer supplemented with RNase A/T1 (1:520, Thermo Fisher Scientific) for 5 min at 22 °C, trimming any exposed RNA ends. The process was stopped by washing each sample with 200 μl of wash buffer supplemented with SUPERase In RNase inhibitor (final concentration of 0.1 U/μL, Ambion) three times for 5 min at 4 °C. Following PNK treatment for 2 h at 22 °C (1x PNK buffer A, 1 mM ATP, 1 U/μl recombinant RNAse inhibitor, 0.5 U/μl T4 PNK (New England BioLabs)), samples were washed again three times with wash buffer-RIn. Hfq-bound RNAs were then proximity ligated with T4 RNA ligase I (1x T4 RNA ligase buffer, 9% DMSO, 1 mM ATP, 20% PEG 8000, 0.6 U/μl recombinant RNase inhibitor, 2.7 U/μl T4 RNA ligase I (New England BioLabs)) overnight at 22°C with agitation, followed by 3 washing steps with wash buffer-RIn at 4 °C. Finally, RNA was eluted by incubating the beads with Proteinase K (50 mM Tris-HCl pH 7.8, 50 mM NaCl, 1% SDS, 5 mM EDTA pH 8.0, 5 mM β-mercaptoethanol, 0.1 U/μl recombinant RNase inhibitor, 0.33 mg/ml Proteinase K (Thermo Fisher Scientific)) for 2 h at 55 °C. RNA purification was achieved using TRIzol LS according to the manufacturer’s instructions. Purified RNA was resuspended in 7 μL of nuclease-free water and quality controlled on a Bioanalyzer Pico RNA chip.

cDNA library preparation and sequencing were performed by Vertis Biotechnologie AG. First, oligonucleotide adapters were ligated to the RNA 5’ and 3’ends followed by first-strand cDNA synthesis using M-MLV reverse transcriptase and the 3’ adapters as primers. The resulting cDNAs were PCR-amplified using a high fidelity DNA polymerase for 14-16 cycles. Next, cDNA was purified using the Agencourt AMPure XP kit (Beckman Coulter Genomics) and quality controlled by capillary electrophoresis on a Shimadzu MultiNA microchip electrophoresis system. For Illumina NextSeq sequencing, the samples were pooled in approximately equimolar amounts. The resulting cDNA pool was then size fractionated in the size range of 170 – 400 bp by polyacrylamide gel electrophoresis and paired-end sequenced on an Illumina HighSeq system using 2 × 150 bp read lengths.

#### RIL-seq data analysis

Sequencing and mapping results are listed in Supplementary Table 1. Data analysis was performed as described in Melamed *et al*., with a few modifications^24,25^. Briefly, raw reads were trimmed and quality filtered with BBDuk (min. phred-score of 20, min. read length of 25). First mapping was conducted with BWA provided by the RIL-seq computational pipeline^72^. The map_chimeric_fragments.py script provided by the RIL-seq pipeline was used to classify fragments into single and chimeric. All parameters were set to default. Fragments that mapped within a distance of 1,000 nt or within the same transcript were considered single, whereas fragments that mapped to two different loci were considered chimeric. To test whether two fragments mapped within the same transcript, an additional annotation file for CP010905.2 was used. To decide if the replicates can be considered reproducible, their correlation was computed as described in Melamed *et al*, by comparing the numbers of mapped fragments in corresponding genomic windows between each pair of libraries, for single and chimeric fragments, respectively^25^. To be able to use the ribozero option of RILseq_significant_regions.py, which excludes rRNAs from the analysis, the necessary BioCyc database was generated with Pathway Tools based on the CP010905.2 annotation^73^. The high correlation coefficient of all *hfq*::3xFLAG pairs (r≥0,79) allowed us to unify the replicates into a single dataset (Supplementary Figure 1A). Fisher’s exact test was applied to assign an odds ratio and a p-value to each chimera. Chimeras with a min. odds ratio of 1 and a p-value < 0.05 were considered significant and termed S-chimeras. In addition, only S-chimeras covered by ≥ 25 chimeric reads were considered for further analysis. The final output table was merged with the CP010905.2 annotation manually curated by assigning each interaction partner to one of the following categories: sRNA, 5’UTR, riboswitch, coding sequence (CDS), 3’UTR, CRISPR, tRNA, intergenic region (IGR), or anti-sense (AS), resulting in Supplementary Table 2 & 3^18^. Pairs can appear more than once if corresponding chimeric reads span multiple regions, such as an mRNA 5’UTR and CDS. Only counting each pair once, the dataset consisted of 1,569 unique interactions (1,198,921 chimeric reads) in the Hfq-FLAG strain and 6 interactions (461 chimeric reads) in the WT, yielding similar numbers to published RIL-seq data (Supplementary Table 1)^24,31^.

To analyse intra-operon RBS overlaps of interactions, an inhouse python script was used. First a database of all intra-operon RBSs was build, based on the operon table published in Fuchs *et al*^18^. The table was filtered for operons with primary TSSs, excluding the first gene of an operon. Intra-operon RBS regions were defined as 25 nt upstream and 20 nt downstream of the respective start codon of the gene. RNA1 and RNA2 of all S-chimeras were searched for overlaps with intra-operon RBS regions. For this purpose, the coordinates listed in the S-chimera table were used whereas the coordinate of either RNA1 start of last read and RNA2 start of first read were extended 100 nt in the respective direction. Additionally, the final output of interactions overlapping with intra-operon RBS regions was curated manually (Supplementary Table 5).

#### RIL-seq data visualization

To count how many chimeric or single reads overlap with given features in the CP010905.2 annotation, the script count_chimeric_reads_per_gene.py of the RILseq computational pipeline was used with slight modifications. We modified the script in a way that it would consider a list of feature categories instead of a single one. The following features were considered for the counting: CDS, 3’UTR, 5’UTR, ncRNA (which includes riboswitches, tmRNA, SRP_RNA, RNase_P_RNA, 6S RNA), sRNAs, rRNA, tRNA, antisense and intergenic regions. A fragment was counted as intergenic if it did not overlap with any of the other features. The minimal overlap between a fragment and a gene was set to 5. If any fragment overlapped with two features, both were counted.

To create images of specific interactions, bed files of chimeric fragments of single interactions were generated using the script generate_BED_file_of_endpoints.py of the RIL-seq computational pipeline. The bed files were visualized with IGV 2.12.3, and further processed with Inkscape 0.92.4 for the respective figure (Figure 2C). For coverage plots these bed files were first converted into coverage files with bedtools genomecov, followed by plot generation using the R package Gviz (*eg*. Figure 3D)^74,75^.

For circos plot visualization of sRNA networks, data were obtained by using the script plot_circos_plot.py from the RIL-seq computational pipeline. Circos plots were generated with Circos^76^. To avoid overloading the plots, only a fraction of the interactions were shown, as indicated in the respective figure description. For a more detailed visualization of all SpoY and SpoX target interactions Cytoscape 3.9.1 was used. All targets (nodes) were included, with targets supported by ≥25 chimeras marked by a solid line, while targets supported by <25 chimeras were highlighted with a dashed line. Target types were discriminated by color as indicated in the figure legend. Edge strengths correlate with the total number of chimeras supporting an individual interaction as listed in Supplementary Table 2.

### Hot phenol extraction of total RNA

Total RNA was extracted using the hot phenol protocol. Bacterial cultures were grown to the desired OD_600_, mixed with 0.2 volumes of STOP solution (95% ethanol, 5% phenol) and snap-frozen in liquid nitrogen. Once thawed on ice, the cell suspension was centrifuged for 20 min, 4500 rpm at 4 °C and the supernatant discarded. For cell lysis, pellets were suspended in 600 μl of 10 mg/ml lysozyme in TE buffer (pH 8.0) and incubated at 37 °C for 10 min. Next, 60 μl of 10% w/v SDS was added respectively, samples were mixed by inversion, and incubated in a water bath at 64 °C, 1-2 min before adding 66 μl 3 M NaOAc, pH 5.2. Phase separation was induced by mixing samples with 750 μl of acid phenol (Roti-Aqua phenol), followed by incubation for 6 min at 64 °C, while regularly inverting the tubes. Samples were briefly placed on ice to cool before centrifugation for 15 min, 13,000 rpm at 4 °C. The aqueous layer was transferred into a 2 ml phase lock gel tube (Eppendorf) and mixed with 750 μl chloroform (Roth) by shaking, followed by centrifugation for 12 min, 13,000 rpm at room temperature. For ethanol precipitation, the aqueous layer was transferred to a new tube, 2 volumes of a 30:1 EtOH:3 M NaOAc, pH 6.5 mix was added and incubated overnight at −20 °C. Finally, samples were centrifuged, washed with cold 75% v/v ethanol and air-dried for 15 min. Precipitated RNA was resuspended in 50 μl nuclease-free water and stored at −80 °C.

### Northern blotting

RNA was purified as described above. Samples were mixed with equal amounts of gel loading buffer II (95% formamide, 18 mM EDTA, 0,025% SDS, 2% bromphenolblue), boiled at 98 °C for 5 min and cooled down on ice before loading on a denaturing 6% polyacrylamide gel containing 7 M urea. RNA was separated for 1 h and 50 min, 300 V and transferred onto a Hybond-N+ membrane (GE Healthcare Life Sciences) at 4 °C for 1 h, 50 V (∼100 W) followed by UV irradiation (0.12 J/cm^2^). Once cross-linked, membranes were pre-hybridized for 10 min in ROTI Hybri-Quick Buffer (Roth) before adding P^32^-labeled DNA oligonucleotides. 5’-labeling was performed by incubating 10 pmol oligonucleotide with 1 μL of ^32^P-γ-ATP (10 μCi/μL) and 5 U T4 Polynucleotide Kinase (Thermo Fisher Scientific) for 1 h at 37°C in a 10 μL reaction. Labeled oligonucleotides were purified using microspin G-25 columns (GE Healthcare) according to the manufacturer’s instructions. Following hybridization overnight at 42 °C, membranes were washed three times with decreasing concentrations (5x, 1x and 0.5x) of SSC buffer (20x SSC: 3 M NaCl, 0.3 M sodium citrate, pH 7.0). Air dried membranes were then exposed onto a phosphor screen for 1-7 days and signals were visualized on a Typhoon FLA 7000 phosphor imager. Following signal detection, the membranes were stripped (0.1% SDS in freshly boiled water, 15 min), and incubated with ROTI Hybri-Quick Buffer (Roth) before adding a new P^32^-labeled DNA oligonucleotide.

### In vitro transcription and radiolabeling of RNA

For *in vitro* transcription of SpoY, SpoX, *spo0A* and associated mutants, pFF-162/-163/-164/-245/-166/-248/-247 and corresponding primers were used for template generation via Phusion High-Fidelity PCR. Resulting PCR products were purified from 1% agarose gels with NucleoSpin Gel and PCR Clean-Up Kit (Macherey-Nagel) to prevent the production of side products during *in vitro* transcription. *In vitro* transcription was performed using the Invitrogen MEGAscript T7 Transcription Kit (Thermofisher Scientific) in 40 μl reactions according to the manufacturer’s protocol. Resulting RNA fragments were separated on a denaturing urea PAGE with 6% polyacrylamide and 7 M urea, followed by ethidium bromide (Carl Roth) staining for 10 min and imaging using an Intas Gel Doc system. Bands of correct size were cut out in small pieces and transferred into 2 ml tubes. For RNA elution, 750 μl RNA elution buffer (0.1 M NaAc, 0.1% SDS, 10 mM EDTA) was added and the samples were incubated at 4 °C and 1000 rpm overnight. Following centrifugation at 5,000 x g and 4 °C for 1 min, the supernatants were transferred to new tubes and RNA extraction was performed using a single phenol-chloroform extraction step (ROTI phenol/chloroform/isoamylalkohol). Purified RNA was resuspended in 20 μl RNase-free water and stored at −80 °C. For radioactive labeling, 50 pmol of *in vitro* transcribed SpoY, SpoY*, SpoX or SpoX* was dephosphorylated using 25 U of calf intestine alkaline phosphatase (NEB) in a 50 μL reaction volume and incubated for 1 h at 37 °C. RNA was extracted again, using a single phenol-chloroform extraction step (ROTI phenol/chloroform/isoamylalkohol) and resuspended in 16 μl RNase-free water. Subsequently, 20 pmol of dephosphorylated and purified RNA was 5′ end-labeled (20 μCi of ^32^P-γATP) using 1 U of Polynucleotide Kinase (NEB) for 1 h at 37 °C in a 20 μL reaction volume. Finally, the labeled RNA was purified on a G-50 column (GE Healthcare) according to manufacturer’s instructions and extracted from a polyacrylamide gel as described above following visualization on a Phosphorimager (FLA-3000 Series, Fuji). Purified RNA was resuspended in 10 μl RNase-free water and stored at −80 °C for up to 2 weeks.

### EMSA (electrophoretic mobility shift assays)

EMSAs were performed by incubating 0.04 pmol of radio-labelled sRNA either alone or with increasing concentrations of *in vitro* transcribed mRNA. Prior to incubation, labeled sRNA and unlabeled mRNA were denatured at 95 °C for 1 min and chilled on ice for 5 min. All components were mixed to a final concentration of 1x structure buffer (10 mM Tris-HCl pH 7.0, 0.1 M KCl, 10 mM MgCl_2_), 0.004 pmol/μl sRNA, 0.1 μg/μl yeast RNA (Ambion) and mRNA ranging from 0-1,000 pmol/μl. For EMSAs analyzing SpoY interactions, 500 pmol/μl purified *C. difficile* Hfq_6_ was added as well ^18^. Reactions were incubated at 37°C for 1 h, stopped by adding 3 μl of 5x native loading dye (0.5x TBE, 50% glycerol, 0.2% xylene cyanol, 0.2% bromophenol blue) and directly loaded on a native 6% polyacrylamide gel at 4 °C in 0.5% TBE at 300 V for 3-4 h. The gel was dried for 1 h at 80 °C on a Gel Dryer 583 (Bio-Rad) and visualized after appropriate exposure on a Phosphorimager (FLA-3000 Series, Fuji).

### In-line probing

In line probing exploits the natural instability of unpaired RNA that leads to differential degradation according to its structure, allowing elucidation of secondary structure information. In-line probing assays were performed by incubating 0.2 pmol of labeled sRNAs either alone or with increasing concentrations of *in vitro* transcribed *spo0A* (0.2, 2, and 20 pmol) for 40 h at room temperature in 1x in-line probing buffer (100 mM KCl, 20 mM MgCl_2_, 50 mM Tris-HCl, pH 8.3). Both, sRNAs and mRNA were denatured at 95 °C for 1 min and chilled on ice for 5 min before assembling the reactions. Ladders were prepared directly prior to loading. For the RNase T1 ladder 0.2 pmol of labeled sRNA was incubated with 8 μL of 1x sequencing buffer (Ambion) at 95 °C for 1 min followed by the addition of 1 μL RNase T1 (0.1 U/ μl) and incubation at 37 °C for 5 min. The alkaline hydrolysis ladder was prepared by incubating 0.2 pmol labeled sRNA with 9 μL of 1x alkaline hydrolysis buffer (Ambion) and incubated at 95 °C for 5 min. All reactions were stopped by the addition of 10 μL of 2x colourless gel-loading solution (10 M urea, 1.5 mM EDTA) and stored on ice. 0.2 pmol of labeled sRNA mixed with 10 μL 2x colourless gel-loading solution served as a control. Samples were resolved on a 10% (vol/vol) polyacrylamide, 7 M urea sequencing gel pre-run for 30 min, 45 W prior to sample loading. The gel was dried for 2 h on a Gel Dryer 583 (Bio-Rad) and visualized after appropriate exposure on a Phosphorimager (FLA-3000 Series, Fuji).

### Reporter system assay

Single colonies of *C. difficile* 630 ΔSpoY harbouring pFF-185/-186/-191/-254 or −285 (FFS-536/-535/-537/-779/-798) and *C. difficile* ΔSpoX harbouring pFF-185/-187/-192/-260 or −289 (FFS-539/-538/-540/-785/-802) were used to inoculate overnight cultures in biological triplicates in sterile filtered TY supplemented with TAP. Main cultures were inoculated by diluting overnight cultures 1:330 in sterile filtered TY supplemented with TAP and grown for 6 h before harvesting 0.07 ODs respectively in a 96-well plate (5 min 4500 x g). Cell pellets were resuspended in 200 μl 4% PFA and incubated at room temperature for 30 min in the dark. Following cell fixation, samples were washed three times in 200 μl 1x PBS, resuspended in 30 μl 1x PBS and incubated overnight at 4 °C in the dark, allowing full maturation of mCherry. Subsequently, samples were diluted in 1x PBS to a final volume of 200 μl and mCherry fluorescence was detected using an Agilent NovoCyte Flow Cytometer. The sample acquisition threshold was set to 5,000, ungated in the FSC-H channel, and a maximum of 100,000 events. Three parameters were recorded for each particle, including FSC-H, SSC-H and PE-Texas Red-H.

For northern blot validation of sRNA expression from reporter system constructs, FFS-535/-536/-537 and FFS-538/-539/-540 were used to inoculate overnight cultures in BHI supplemented with TAP in biological duplicates. Main cultures were inoculated to an OD_600_ of 0.05 and grown to a final OD_600_ of 1 before harvesting cells and extracting RNA using the hot phenol protocol described above. Northern blotting was performed as described above and sRNA expression could be confirmed, as illustrated in Supplementary Figure 5E&F

### Western blotting

To test Spo0A expression in a SpoY and SpoX deletion mutant and corresponding overexpression strains, FFS-591, FFS-593, FFS-535, FFS-594 and FFS-538 were inoculated into sterile filtered TY supplemented with TAP in biological duplicates from single colonies. Main cultures were inoculated by diluting overnight cultures to OD_600_ = 0.05 in sterile filtered TY containing TAP. For western blot analysis samples were taken at mid-exponential (5.5 h post inoculation) and stationary phase (9 h post inoculation) by harvesting 2 OD units *via* centrifugation for 5 min at 5,000 x g. Pellets were frozen overnight at −20°C, followed by cell resuspension in 50 μl 1x PBS and incubation for 50 mins at 37°C, leading to cell lysis^77^. Subsequently, cell lysates were mixed with equal amounts of 2x protein loading dye and boiled for 5 min at 95 °C. Of each sample, 0.3 OD (15 μl) was loaded and separated on a 15% SDS–polyacrylamide gel followed by transfer of proteins to a Protran 0.2 μm NC membrane (Amersham) at 4 °C for 1.5 h, 340 mM using a semi-dry blotting system. Equal loading of protein samples was confirmed *via* Ponceau S staining (Sigma-Aldrich) for 4 min. Staining was reversed by washing the stained membrane with 0.1 M NaOH for 1 min, followed by blocking in TBS-T with 5% powdered milk for 1 h at room temperature. Subsequently the membrane was incubated overnight at 4 °C with anti-Spo0A antibody diluted 1:5,000 in TBS-T with 5% powdered milk and washed again 3x in TBS-T for 10 min. Following the last washing step the membrane was incubated for 1 h at room temperature with anti-mouse-HRP antibody (Thermo Scientific) diluted 1:10,000 in TBS-T with 5% powdered milk and finally washed 3x in TBS-T for 10 min before adding ECL substrate (Amersham) for detection of HRP activity using a CCD camera (ImageQuant, GE Healthcare).

### RT-qPCR analysis of sporulation genes

Glycerol stocks of *C. difficile* strains (FFS-535, FFS-538, FFS-591, FFS-593, and FFS-594) were inoculated onto BHIS plates (BHI agar containing 5 g/L yeast extract (Roth) and 0.1% sterile filtered cysteine) supplemented with taurocholate (TA, 0.1% w/v) and TAP. Single colonies were then inoculated into liquid BHIS-TA-TAP media in biological triplicates and grown overnight. These cultures were diluted to OD_600_ = 0.05 with BHIS-TA-TAP media and grown till early stationary phase. Pre-cultures were then diluted using BHIS-TAP to OD_600_ = 0.05 and grown as main-cultures until an OD_600_ of ∼0.5. To induce sporulation, 120 μl of the main cultures were spread onto 70:30 agar plates (70% SMC media and 30% BHIS media) supplemented with TAP. Sporulating cultures were collected at 9 h and 12 h after plating using phosphate-buffered saline (PBS) and flash frozen immediately upon the addition of 0.2 volumes ice-cold STOP mix (5% water-saturated phenol (pH < 7.0) in ethanol). Total RNA was then isolated from collected samples using the hot-phenol extraction procedure as described above. For DNA removal, 5 μg of total RNA was treated with DNase I (Thermo Scientific) for 1 h at 37 °C and then further purified using a single phenol-chloroform extraction step (ROTI Phenol/Chloroform/Isoamylalkohol). Purified RNA was resuspended in 50 μl RNase-free water and stored at −80 °C. Reverse transcription was performed using the M-MLV Reverse Transcriptase Kit (Invitrogen) as per manufacturer’s instructions, with 1 μg of DNase I treated, purified total RNA and Random Hexamer Primer (Invitrogen). cDNA was then diluted 20-fold and 1 μL were used for each qPCR reaction along with 20 nM of gene-specific oligonucleotides in a 10 μL reaction mix. qPCR was performed with Takyon™ No ROX SYBR 2x MasterMix blue dTTP (Eurogentec) reagent in technical duplicates using the QuantStudio™ 5 Real-Time PCR cycler (Thermo Fisher Scientific) and following conditions: TakyonTM activation at 95 °C for 3 min; DNA denaturing at 95 °C for 10 sec; 40 cycles of annealing and extension at 60 °C for 60 sec, followed by melting curve denaturation at 95 °C for 15 sec and melting curve analysis at 55-95 °C using “step and hold” with 0.5 °C and 10 sec of incubation per step. Transcript levels were normalized to 5S rRNA and are displayed as Ct values, representing Log_2_-fold change relative to FFS-591 (WT-*p*[ctl]). All oligonucleotide sequences used for RT-qPCR are listed in Supplementary Table 7.

### Sporulation frequencies

Sporulation assays were performed in 70:30 sporulation broth medium according to Edwards *et al* with minor alterations^78^. In short, *C. difficile* cultures (FFS-535, FFS-538, FFS-591, FFS-593, and FFS-594) were started in biological triplicates in BHIS medium supplemented with 0.1% taurocholate (TA) and 0.2% fructose until mid-log phase (OD_600_ ≤ 0.9). Cultures were then back-diluted in 70:30 medium to OD_600_ = 0.01 and monitored for the production of spores. At each timepoint (6 h, 12 h, 24 h and 48 h post inoculation), samples were taken, serially diluted, and spotted (10 μl spots in technical triplicates) on BHIS-TA plates and incubated for 24 to 48 h to enumerate total number of CFU (spores and vegetative cells). Simultaneously, 500 μl from each culture was removed, mixed 1:1 with 95% EtOH and incubated for 30 min to kill all vegetative cells. EtOH-treated samples were then serially diluted, similarly plated, and incubated, representing the spore CFU. The sporulation frequency was determined by dividing the number of spores by the total number of CFUs at each time point (spore ratio), multiplied by 100 (percentage of spores formed).

For phase-contrast microscopy *C. difficile* strains were grown in 70:30 sporulation medium as described above in biological triplicates. At 12 h, 24 h and 48 h post inoculation, 1 ml of culture was removed from the anaerobic chamber, centrifuged at full speed for 30 s, and resuspended in ∼10-30 μl of supernatant. Microcopy slides were prepared by placing 2 μl of the concentrated cultures onto thin 1% agarose pads that were applied directly to the surface of the slide. Phase-contrast microscopy was performed using a HC PLAN FLUOTAR 100x/1.32 PH3 oil immersion objective on a LEICA DM2500 microscope. Two fields of view for each strain and replicate were acquired and used to calculate the percentage of spores (the number of spores divided by the total number of spores, prespores, and vegetative cells; 300 cells per field of view were analyzed).

### Murine model of C. difficile infection

All animal experiments were performed in agreement with the guidelines of the Helmholtz Centre for Infection Research (HZI), Brunswick, Germany, the national animal protection law (TierSchG), the animal experiment regulations (TierSchVersV), and the recommendations of the Federation of European Laboratory Animal Science Association (FELASA). Mice experiments were approved by the Lower Saxony State Office for Nature, Environment and Consumer Protection (LAVES), Oldenburg, Lower Saxony, Germany; permit No. 33.19-42502-04-19/3126.

C57BL/6N SPF mice were maintained (including housing) at the animal facilities of the HZI under enhanced specific pathogen-free (SPF) conditions for at least two weeks before the start of the experiment. Female mice aged between 12-14 weeks were used. Sterilized food and water were provided *ad libitum*. Mice were kept under a strict 12-hour light cycle (lights on at 7:00 am and off at 7:00 pm) and housed in groups of up to six mice per cage. All mice were euthanized by asphyxiation with CO_2_ and cervical dislocation.

Infection experiments were performed with *C. difficile* 630 strain FFS-01 (WT), FFS-491 (ΔSpoY) and FFS-492 (ΔSpoX). SPF mice were weighted and treated with 10 mg/kg clindamycin 24 h prior to infection, administered via intraperitoneal injection to induce susceptibility to *C. difficile* infection^79^. Spores were heat-treated at 65 °C for 20 min before infection to kill remaining vegetative cells. Mice were infected with 10^4^ *C. difficile* spores in 200 μL 1x PBS administered *via* oral gavage. Following infection, mice were monitored and scored daily for symptoms of clinically severe CDI including behavior, posture, fur and skin, provoked behavior, weight loss and feces consistency. Mice showing signs of CDI were monitored twice a day and euthanized after losing 20% of their initial weight or developing severe clinical signs of features listed above.

For quantification of bacterial burden, fresh fecal samples were collected at different time points, their weight recorded, supplemented with 1.0 mm diameter zirconium/glass beads in 1 mL 1x PBS and subsequently homogenized for 50 sec with Mini-Beadbeater-96 (Biospec). To determine colony forming units (CFUs), serial dilutions of homogenized samples were plated on bioMérieuxTM *C. difficile* agar. For the quantification of spores, aliquots of homogenized samples were incubated 65 °C for 20 min to kill remaining vegetative cells and plated on bioMérieux™ *C. difficile* agar pretreated with 0.1% taurocholic acid to induce germination. Plates were cultured at 37 °C for 48 h in anaerobic jars before counting. CFUs of *C. difficile* were calculated by normalization to feces weight.

### Prediction of RNA folding and sRNA target interactions

Secondary structures of sRNAs were predicted with the RNAfold WebServer, while RNAcofold was used to predict secondary structures of single stranded sRNA and mRNA sequences upon dimer formation^46,80^. In both cases, structures were visualized with VARNA^81^.

Potential sRNA-target interactions were predicted using IntaRNA by uploading either SpoY, SpoX or *spo0A* in combination with interaction partners revealed by RIL-seq analysis (Supplementary Table 2)^47^. Default settings were used.

### Prediction of sRNA target motif

Motif search and generation of sequence logos was accomplished with MEME version 5.4.1^45^. For both sRNAs, all target sequences were extracted from the RIL-seq data, including target interactions supported by <25 chimeras (Supplementary Table 2). For this purpose, the coordinates listed in the S-chimera table for “start of first read” and “start of last read” were used and extended 50 nt downstream. The resulting sequences were uploaded to MEME for target motif identification (SpoY = 28 targets, SpoX = 42 targets). The number of motifs to be found by MEME was set to 5 and only the given strand was searched. All other settings were left at default.

### Quantification and Statistical analysis

Statistical analysis of RIL-seq results is described above and was performed according to the original protocol^24,25^. Quantification and analysis of western blot, northern blot and EMSA signals as well as in-line probing and microscopy images was performed with ImageJ^82^. GraphPad Prism 9 was used for all statistical analyses and data visualizations, in combination with Inkscape 0.92.4^82^. Sample sizes and detailed descriptions of statistical analyses are indicated in the figure legends and method section for each experiment separately.

## DATA AVAILABILITY

All RNA-sequencing data are available at the National Center for Biotechnology Information Gene Expression Omnibus database (https://www.ncbi.nlm.nih.gov/geo) under the accession number GSE213005. The RIL-seq dataset can be accessed *via* an RNA-RNA interactome browser (available upon publication).

## ACKNOWLEDGEMENT

We would like to thank Gianluca Matera for assisting with the RIL-seq experiment and Aimee Shen for providing anti-Spo0A antibodies. We are also very grateful to Anke Sparmann and Jörg Vogel for providing critical and insightful feedback on the manuscript. T.L. was supported by bayresq.net. J.S. was supported by a DFG grant (FA 1113/2-1). M.G. was supported by the European Social Fund (ESF) under Grant 823905.

## AUTHOR CONTRIBUTIONS

M.F. co-designed all experimental work, performed the majority of the experimental work, and co-wrote the manuscript. V.L.-S. performed all bioinformatic analyses. J.S. and T.L. performed parts of the experimental work. M.G. setup the RIL-seq browser. A.B. and T.S. co-designed and performed mouse experiments. All co-authors provided feedback on the manuscript. F.F. co-designed all experiments and co-wrote the manuscript.

## REFERENCES

1. Centers for Disease Control and Prevention. Antibiotic resistance threats in the United States, 2019. CDC (2019).

2. European Centre for Disease Prevention and Control. Healthcare-associated infections: Clostridium difficile infections. ECDC (2018).

3. Bartlett, J. G., Chang, T. W., Gurwith, M., Gorbach, S. L. & Onderdonk, A. B. Antibiotic-Associated Pseudomembranous Colitis Due to Toxin-Producing Clostridia. N. Engl. J. Med. 298, 531–534 (1978).

4. Smits, W. K., Lyras, D., Lacy, D. B., Wilcox, M. H. & Kuijper, E. J. Clostridium difficile infection. Nat. Rev. Dis. Prim. 2, 16020 (2016).

5. Peng, Z. et al. Update on antimicrobial resistance in Clostridium difficile: resistance mechanisms and antimicrobial susceptibility testing. J. Clin. Microbiol. 55, 1998–2008 (2017).

6. O’Grady, K., Knight, D. R. & Riley, T. V. Antimicrobial resistance in Clostridioides difficile. Eur. J. Clin. Microbiol. Infect. Dis. (2021) doi:10.1007/s10096-021-04311-5.

7. Zhu, D., Sorg, J. A. & Sun, X. Clostridioides difficile biology: sporulation, germination, and corresponding therapies for C. difficile infection. Front. Cell. Infect. Microbiol. 8, 29 (2018).

8. Edwards, A. N. & McBride, S. M. Initiation of sporulation in Clostridium difficile: a twist on the classic model. FEMS Microbiol. Lett. 358, 110–118 (2014).

9. Shen, A., Edwards, A. N., Sarker, M. R. & Paredes-Sabja, D. Sporulation and germination in clostridial pathogens. in Gram-Positive Pathogens, Third Edition vol. 7 903–926 (American Society of Microbiology, 2019).

10. Deakin, L. J. et al. The Clostridium difficile spo0A gene is a persistence and transmission factor. Infect. Immun. 80, 2704–2711 (2012).

11. Rosenbusch, K. E., Bakker, D., Kuijper, E. J. & Smits, W. K. C. difficile 630Δerm Spo0A regulates sporulation, but does not contribute to toxin production, by direct high-affinity binding to target DNA. PLoS One 7, e48608 (2012).

12. Lee, C. D. et al. Genetic mechanisms governing sporulation initiation in Clostridioides difficile. Curr. Opin. Microbiol. 66, 32–38 (2022).

13. Shen, A., Edwards, A. N., Sarker, M. R. & Paredes-Sabja, D. Sporulation and germination in Clostridial pathogens. in Gram-Positive Pathogens, Third Edition vol. 7 903–926 (American Society of Microbiology, 2019).

14. Boudry, P. et al. Pleiotropic role of the RNA chaperone protein Hfq in the human pathogen Clostridium difficile. J. Bacteriol. 196, 3234–3248 (2014).

15. Maikova, A., Kreis, V., Boutserin, A., Severinov, K. & Soutourina, O. Using an endogenous CRISPR-Cas system for genome editing in the human pathogen Clostridium difficile. Appl. Environ. Microbiol. 85, (2019).

16. Holmqvist, E. & Vogel, J. RNA-binding proteins in bacteria. Nat. Rev. Microbiol. (2018) doi:10.1038/s41579-018-0049-5.

17. Boudry, P. et al. Identification of RNAs bound by Hfq reveals widespread RNA partners and a sporulation regulator in the human pathogen Clostridioides difficile. RNA Biol. 1–22 (2021) doi:10.1080/15476286.2021.1882180.

18. Fuchs, M. et al. An RNA-centric global view of Clostridioides difficile reveals broad activity of Hfq in a clinically important gram-positive bacterium. Proc. Natl. Acad. Sci. 118, (2021).

19. Marchais, A., Duperrier, S., Durand, S., Gautheret, D. & Stragier, P. CsfG, a sporulation-specific, small non-coding RNA highly conserved in endospore formers. RNA Biol. 8, 358– 364 (2011).

20. Silvaggi, J. M., Perkins, J. B. & Losick, R. Genes for small, noncoding RNAs under sporulation control in Bacillus subtilis. J. Bacteriol. 188, 532–541 (2006).

21. Schmalisch, M. et al. Small genes under sporulation control in the Bacillus subtilis genome. J. Bacteriol. 192, 5402–5412 (2010).

22. Lamm-Schmidt, V. et al. Grad-seq identifies KhpB as a global RNA-binding protein in Clostridioides difficile that regulates toxin production. microLife 2, 1–21 (2021).

23. Hör, J., Gorski, S. A. & Vogel, J. Bacterial RNA biology on a genome scale. Mol. Cell 70, 785– 799 (2018).

24. Melamed, S. et al. Global mapping of small RNA-target interactions in bacteria. Mol. Cell 63, 884–897 (2016).

25. Melamed, S. et al. Mapping the small RNA interactome in bacteria using RIL-seq. Nat. Protoc. 13, 1–33 (2018).

26. Helwak, A., Kudla, G., Dudnakova, T. & Tollervey, D. Mapping the human miRNA interactome by CLASH reveals frequent noncanonical binding. Cell 153, 654–665 (2013).

27. Kwok, C. K. Dawn of the in vivo RNA structurome and interactome. Biochem. Soc. Trans. 44, 1395–1410 (2016).

28. Saujet, L., Monot, M., Dupuy, B., Soutourina, O. & Martin-Verstraete, I. The key sigma factor of transition phase, SigH, controls sporulation, metabolism, and virulence factor expression in Clostridium difficile. J. Bacteriol. 193, 3186–96 (2011).

29. Hofmann, J. D. et al. Metabolic reprogramming of Clostridioides difficile during the stationary phase with the induction of toxin production. Front. Microbiol. 9, 1970 (2018).

30. Park, S. et al. Dynamic interactions between the RNA chaperone Hfq, small regulatory RNAs and mRNAs in live bacterial cells. Elife 10, (2021).

31. Matera, G. et al. Global RNA interactome of Salmonella discovers a 5′ UTR sponge for the MicF small RNA that connects membrane permeability to transport capacity. Mol. Cell (2022) doi:10.1016/j.molcel.2021.12.030.

32. Menendez-Gil, P. & Toledo-Arana, A. Bacterial 3′UTRs: a useful resource in post-transcriptional regulation. Front. Mol. Biosci. 7, 617633 (2021).

33. Bronesky, D. et al. A multifaceted small RNA modulates gene expression upon glucose limitation in Staphylococcus aureus. EMBO J. 38, (2019).

34. El Mouali, Y. et al. CRP-cAMP mediates silencing of Salmonella virulence at the post-transcriptional level. PLOS Genet. 14, e1007401 (2018).

35. Ruiz de los Mozos, I. et al. Base pairing interaction between 5′- and 3′-UTRs controls icaR mRNA translation in Staphylococcus aureus. PLoS Genet. 9, e1004001 (2013).

36. Bar, A., Argaman, L., Altuvia, Y. & Margalit, H. Prediction of novel bacterial small RNAs from RIL-seq RNA–RNA interaction data. Front. Microbiol. 12, 635070 (2021).

37. Chen, Y., Indurthi, D. C., Jones, S. W. & Papoutsakis, E. T. Small RNAs in the genus clostridium. MBio 2, e00340–10 (2011).

38. Soutourina, O. A. et al. Genome-wide identification of regulatory RNAs in the human pathogen Clostridium difficile. PLoS Genet. 9, e1003493 (2013).

39. Małecka, E. M., Sobańska, D. & Olejniczak, M. Bacterial chaperone protein Hfq facilitates the annealing of sponge RNAs to small regulatory RNAs. J. Mol. Biol. 167291 (2021) doi:10.1016/j.jmb.2021.167291.

40. Balasubramanian, D. & Vanderpool, C. K. New developments in post-transcriptional regulation of operons by small RNAs. RNA Biol. 10, 337–341 (2013).

41. Rice, J. B., Balasubramanian, D. & Vanderpool, C. K. Small RNA binding-site multiplicity involved in translational regulation of a polycistronic mRNA. Proc. Natl. Acad. Sci. 109, E2691–E2698 (2012).

42. Møller, T., Franch, T., Udesen, C., Gerdes, K. & Valentin-Hansen, P. Spot 42 RNA mediates discoordinate expression of the E. coli galactose operon. Genes Dev. 16, 1696–1706 (2002).

43. Desnoyers, G., Morissette, A., Prévost, K. & Massé, E. Small RNA-induced differential degradation of the polycistronic mRNA iscRSUA. EMBO J. 28, 1551–1561 (2009).

44. Saujet, L., Pereira, F. C., Henriques, A. O. & Martin-Verstraete, I. The regulatory network controlling spore formation in Clostridium difficile. FEMS Microbiol. Lett. 358, 1–10 (2014).

45. Bailey, T. L. & Elkan, C. Fitting a mixture model by expectation maximization to discover motifs in biopolymers. Proceedings. Int. Conf. Intell. Syst. Mol. Biol. 2, 28–36 (1994).

46. Bernhart, S. H. et al. Partition function and base pairing probabilities of RNA heterodimers. Algorithms Mol. Biol. 1, 3 (2006).

47. Mann, M., Wright, P. R. & Backofen, R. IntaRNA 2.0: enhanced and customizable prediction of RNA–RNA interactions. Nucleic Acids Res. 45, W435–W439 (2017).

48. Fimlaid, K. A. et al. Global analysis of the sporulation pathway of Clostridium difficile. PLoS Genet. 9, e1003660 (2013).

49. Westermann, A. J. Regulatory RNAs in virulence and host-microbe interactions. in Regulating with RNA in Bacteria and Archaea vol. 6 305–337 (American Society of Microbiology, 2019).

50. Errington, J. Regulation of endospore formation in Bacillus subtilis. Nat. Rev. Microbiol. 1, 117–126 (2003).

51. Stephenson, K. & Hoch, J. A. Evolution of signalling in the sporulation phosphorelay. Mol. Microbiol. 46, 297–304 (2002).

52. Childress, K. O. et al. The phosphotransfer protein CD1492 represses sporulation initiation in Clostridium difficile. Infect. Immun. 84, 3434–3444 (2016).

53. Edwards, A. N., Wetzel, D., DiCandia, M. A. & McBride, S. M. Three orphan histidine kinases inhibit Clostridioides difficile sporulation. J. Bacteriol. e0010622 (2022) doi:10.1128/jb.00106-22.

54. Ul Haq, I., Brantl, S. & Müller, P. A new role for SR1 from Bacillus subtilis: regulation of sporulation by inhibition of kinA translation. Nucleic Acids Res. 49, 10589–10603 (2021).

55. Ohtani, K. et al. Unique regulatory mechanism of sporulation and enterotoxin production in Clostridium perfringens. J. Bacteriol. 195, 2931–6 (2013).

56. Kim, W. & Lee, Y. Mechanism for coordinate regulation of rpoS by sRNA-sRNA interaction in Escherichia coli. RNA Biol. 17, 176–187 (2020).

57. De Lay, N. & Gottesman, S. A complex network of small non-coding RNAs regulate motility in Escherichia coli. Mol. Microbiol. 86, 524–538 (2012).

58. Mika, F. & Hengge, R. Small RNAs in the control of RpoS, CsgD, and biofilm architecture of Escherichia coli. RNA Biol. 11, 494–507 (2014).

59. Underwood, S. et al. Characterization of the sporulation initiation pathway of Clostridium difficile and its role in toxin production. J. Bacteriol. 191, 7296–7305 (2009).

60. Dawson, L. F., Valiente, E., Faulds-Pain, A., Donahue, E. H. & Wren, B. W. Characterisation of Clostridium difficile biofilm formation, a role for Spo0A. PLoS One 7, e50527 (2012).

61. Taggart, M. G. et al. Biofilm regulation in Clostridioides difficile: novel systems linked to hypervirulence. PLOS Pathog. 17, e1009817 (2021).

62. Denham, E. L. The Sponge RNAs of bacteria –how to find them and their role in regulating the post-transcriptional network. Biochim. Biophys. Acta - Gene Regul. Mech. 1863, 194565 (2020).

63. Iosub, I. A. et al. Hfq CLASH uncovers sRNA-target interaction networks linked to nutrient availability adaptation. Elife 9, (2020).

64. Bradshaw, W. J., Kirby, J. M., Roberts, A. K., Shone, C. C. & Acharya, K. R. Cwp2 from Clostridium difficile exhibits an extended three domain fold and cell adhesion in vitro. FEBS J. 284, 2886–2898 (2017).

65. Reynolds, C. B., Emerson, J. E., de la Riva, L., Fagan, R. P. & Fairweather, N. F. The Clostridium difficile cell wallprotein CwpV is antigenically variable between strains, but exhibits conserved aggregation-promoting function. PLoS Pathog. 7, e1002024 (2011).

66. Mondhe, M., Chessher, A., Goh, S., Good, L. & Stach, J. E. M. Species-Selective Killing of Bacteria by Antimicrobial Peptide-PNAs. PLoS One 9, e89082 (2014).

67. Sully, E. K. & Geller, B. L. Antisense antimicrobial therapeutics. Curr. Opin. Microbiol. 33, 47–55 (2016).

68. Green, M. R. & Sambrook, J. Molecular Cloning: A Laboratory Manual. (Cold Spring Harbor Laboratory Press,U.S., 2012).

69. Ransom, E. M., Ellermeier, C. D. & Weiss, D. S. Use of mCherry red fluorescent protein for studies of protein localization and gene expression in Clostridium difficile. Appl. Environ. Microbiol. 81, 1652–1660 (2015).

70. Kirk, J. A. & Fagan, R. P. Heat shock increases conjugation efficiency in Clostridium difficile. Anaerobe 42, 1–5 (2016).

71. Cartman, S. T., Kelly, M. L., Heeg, D., Heap, J. T. & Minton, N. P. Precise manipulation of the Clostridium difficile chromosome reveals a lack of association between the tcdC genotype and toxin p roduction. Appl. Environ. Microbiol. 78, 4683–4690 (2012).

72. Li, H. Aligning sequence reads, clone sequences and assembly contigs with BWA-MEM. Preprint at https://arxiv.org/abs/1303.3997 (2013).

73. Karp, P. D. et al. Pathway Tools version 23.0 update: software for pathway/genome informatics and systems biology. Brief. Bioinform. 22, 109–126 (2021).

74. Hahne, F. & Ivanek, R. Visualizing genomic data using Gviz and Bioconductor. in Methods in molecular biology (Clifton, N.J.) vol. 1418 335–351 (2016).

75. Quinlan, A. R. BEDTools: the Swiss-army tool for genome feature analysis. Curr. Protoc. Bioinforma. 47, 11.12.1-34 (2014).

76. Krzywinski, M. et al. Circos: An information aesthetic for comparative genomics. Genome Res. 19, 1639–1645 (2009).

77. Fagan, R. P. & Fairweather, N. F. Clostridium difficile has two parallel and essential Sec secretion systems. J. Biol. Chem. 286, 27483–27493 (2011).

78. Edwards, A. N., Nawrocki, K. L. & McBride, S. M. Conserved oligopeptide permeases modulate sporulation initiation in Clostridium difficile. Infect. Immun. 82, 4276–4291 (2014).

79. Theriot, C. M., Bowman, A. A. & Young, V. B. Antibiotic-induced alterations of the gut microbiota alter secondary bile acid production and allow for Clostridium difficile spore germination and outgrowth in the large intestine. mSphere 1, (2016).

80. Lorenz, R. et al. ViennaRNA package 2.0. Algorithms Mol. Biol. 6, 26 (2011).

81. Darty, K., Denise, A. & Ponty, Y. VARNA: Interactive drawing and editing of the RNA secondary structure. Bioinformatics 25, 1974–1975 (2009).

82. Schneider, C. A., Rasband, W. S. & Eliceiri, K. W. NIH Image to ImageJ: 25 years of image analysis. Nat. Methods 9, 671–675 (2012).

