## Supplementary Information for "A network of small RNAs regulates sporulation initiation in *C. difficile*"

This PDF file includes:

- Supplementary Figures
- Tables description for Supplementary Table 1 to 8
- Supplementary Table 6 to 8
- SI References

### SUPPLEMENTARY FIGURES

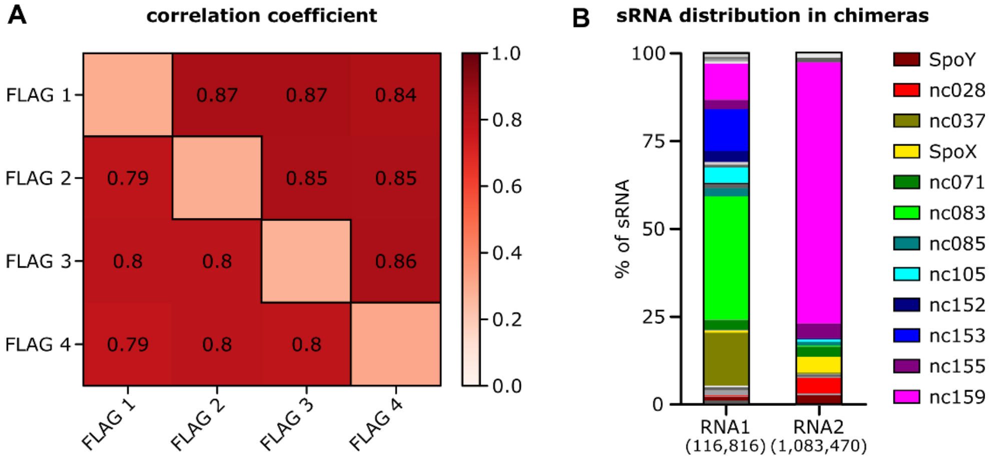

**Supplementary Figure 1: Hfq serves as a platform for RNA-RNA interactions in *C. difficile*. (A)** Replicate reproducibility calculated as correlation coefficient by comparing the numbers of mapped fragments in corresponding genomic windows between each pair of libraries, for single (below diagonal) and chimeric (above diagonal) fragments, respectively. **(B)** Distribution of sRNAs in chimeric fragments, where RNA1 constitutes the 5'end and RNA2 the 3'end of a chimera (n=4). sRNAs that are present in ≥ 1.5% off all chimeras in either RNA1 or RNA2 are highlighted in order of genomic location. A total of 118,740 chimeric reads mapped to sRNAs in RNA1 and 1,091,528 to sRNAs in RNA2**.**

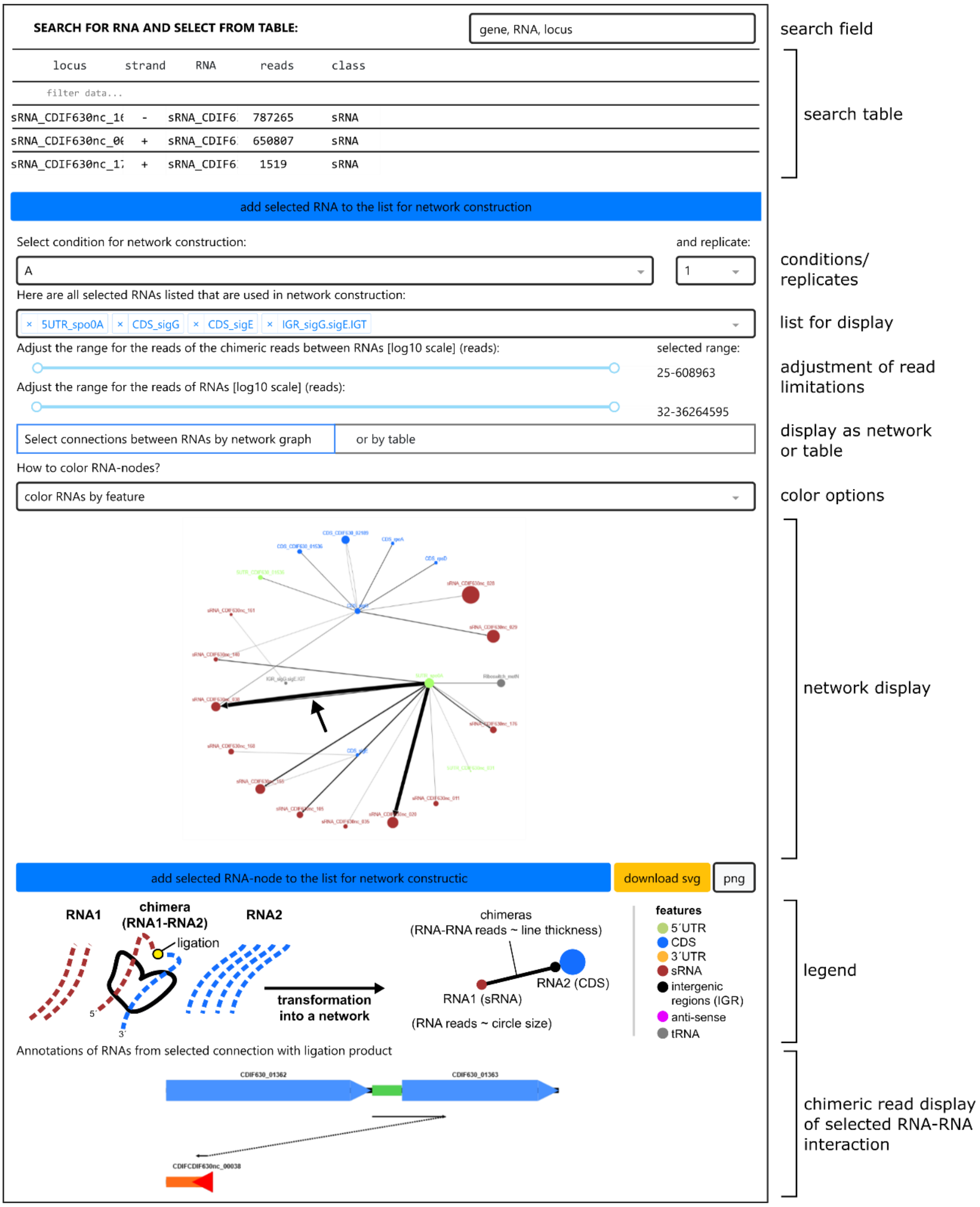

**Supplementary Figure 2: Rilseqcd is a web-browser that allows easy access to our RIL-seq data and an interactive search for RNA-RNA interactions.** Screenshot of the RIL-seq browser accessible *via* https://resources.helmholtz-hiri.de/rilseqcd/. Details explaining the available options are given on the right. So far only one condition and one replicate are available, the latter because all four replicates have been pooled into a single dataset. Specific targets can be search and added to the network display either *via* the search bar and table at the top, or by directly typing into the "list for display" field. If no targets are select, a network of all detected interactions will be shown. By clicking on specific interaction in the "network display", a schematic representation of the selected RNA-RNA interaction will appear on the bottom.

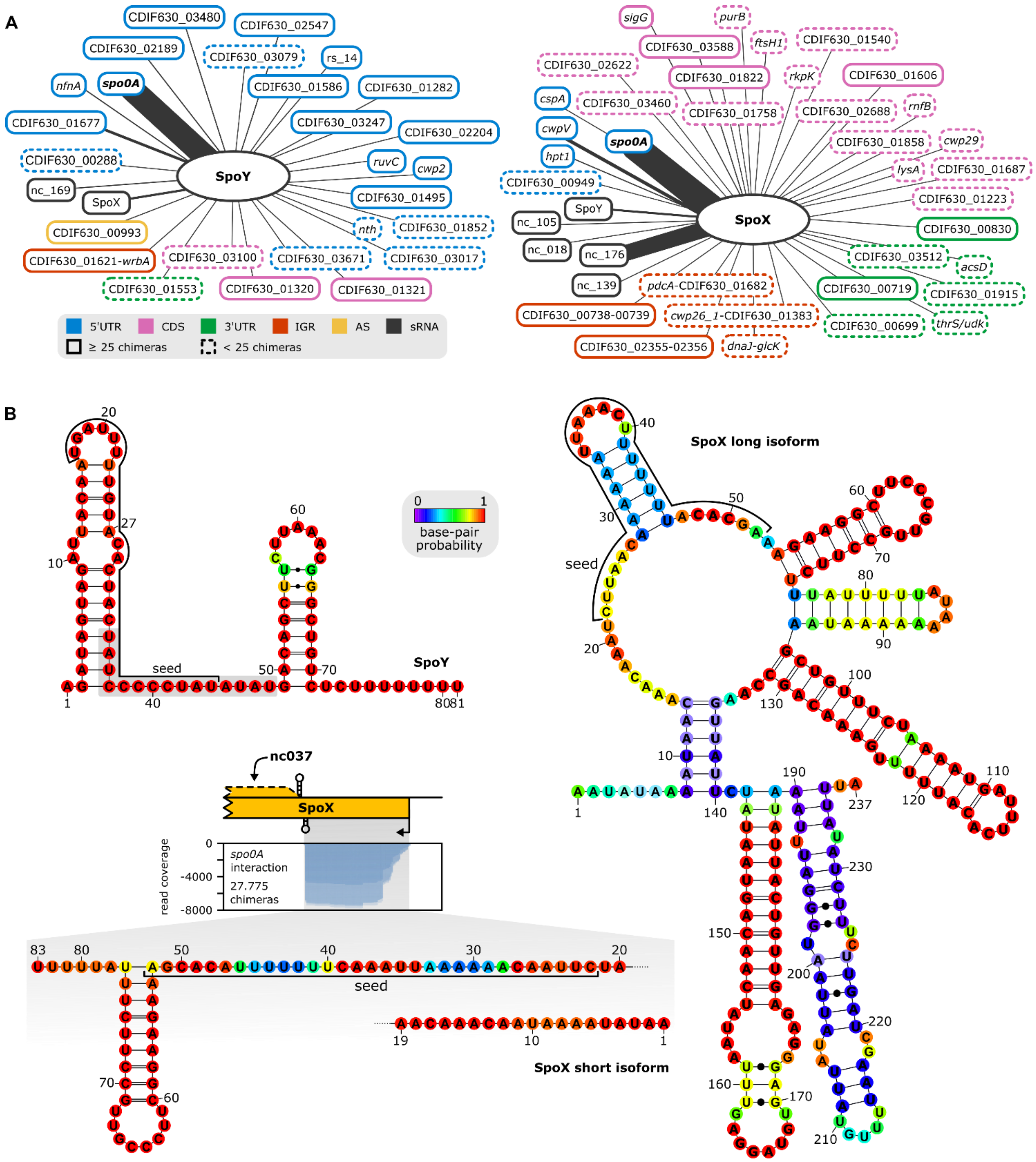

**Supplementary Figure 3: RIL-seq reveals *spo0A* as a target of sRNA-mediated post-transcriptional regulation. (A)** Target network of SpoY and SpoX, targets supported by ≥25 chimeras are marked by a solid line, while targets supported by <25 chimeras are highlighted with a dashed line. Target types are discriminated by color. Edge strength correlates with the number of chimeras supporting an individual interaction. **(B)** Predicted secondary structure (RNAfold[1]) for SpoY and both isoforms of SpoX are provided. Seed regions relevant for *spo0A* interaction were predicted *in silico* (IntaRNA[2]) and emphasized in the secondary structure. Read coverage of SpoX by SpoX-*spo0A* chimeric reads in highlighted in relation to the SpoX encoding region.

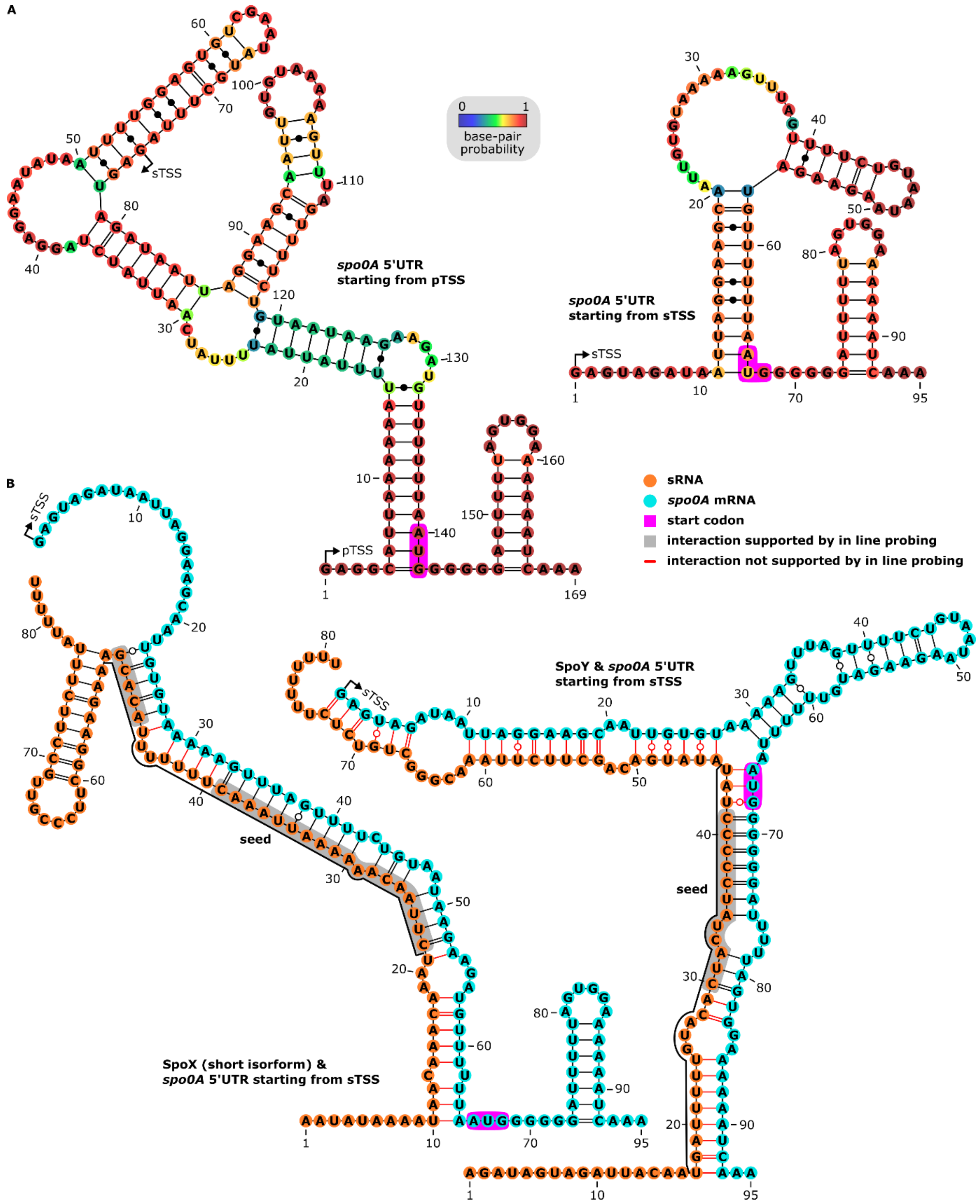

**Supplementary Figure 4: SpoX and SpoY bind *spo0A* at different interaction sites. (A)** Predicted secondary structures of the *spo0A* 5’UTR and beginning of CDS, starting from the primary transcription start site (pTSS) and secondary TSS (RNAfold[1]). The start codon is highlighted in pink. **(B)** Predicted secondary structures of the *spo0A* 5’UTR and beginning of CDS, starting from the sTSS, upon dimer formation with either SpoX (short isoform) or SpoY (RNAcofold[3]). sRNAs and mRNA are highlighted in orange and blue, respectively. sRNA-target base-pairings supported by in line probing (Figure 4B) are shaded in grey, while base-pairings that are not confirmed experimentally are marked in red.

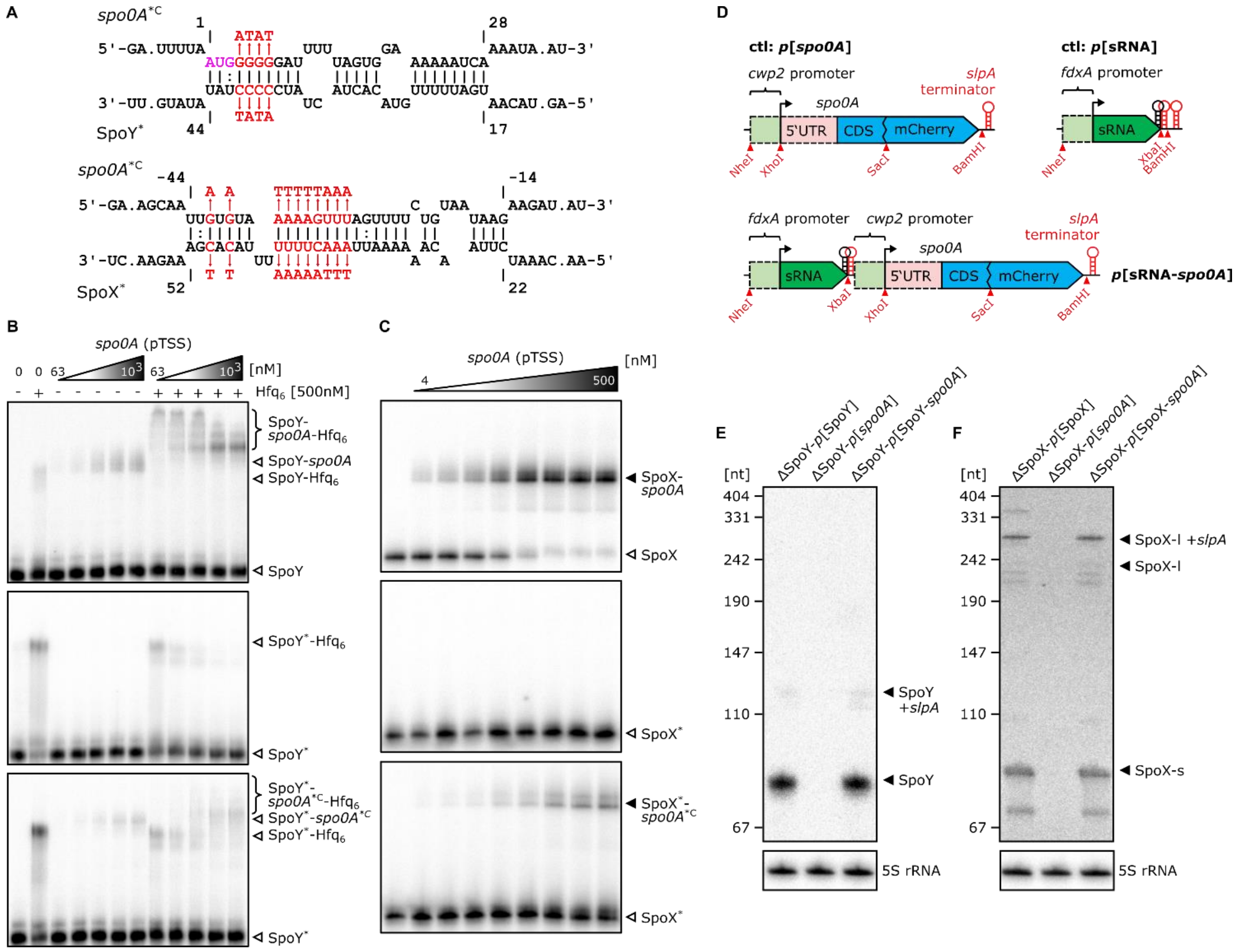

**Supplementary Figure 5**: **SpoY and SpoX directly interact with the *spo0A* mRNA *in vitro* and *in vivo*. (A)** *In silico* predicted SpoY-*spo0A* and SpoX-*spo0A* interaction sites (IntaRNA[2]). Mutations introduced in the sRNA seed region as well as compensatory mutations in the *spo0A* target region are highlighted in red. The *spo0A* nucleotide position is calculated relative to the *spo0A* start codon (highlighted in pink). **(B-C)** EMSAs performed with either $^{32}$P-labeled SpoY (B) or SpoX (short isoform) (C) with increasing concentrations of the *spo0A* target region, respectively. Purified Hfq was added to facilitate SpoY-*spo0A* complex formation. Mutating the respective sRNA seed region (SpoY*/SpoX*) abolished the interaction, while introducing compensatory mutations into the *spo0A* target region (*spo0A**C) slightly rescued the complex formation. A representative image of three independent experiments is shown, respectively. **(D)** Schematic representation of translational fusion constructs designed for *in vivo* reporter system assays. Restriction sites allowing easy exchange of each component individually are annotated. **(E-F)** Northern blot validation of sRNA expression from reporter constructs in the respective sRNA deletion mutant. A representative image of two independent experiments is shown, respectively.

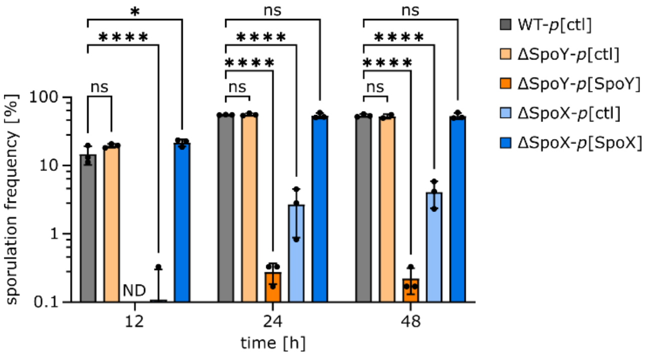

**Supplementary Figure 6: sRNA mediated regulation of *spo0A* affects sporulation frequencies.** Sporulation frequencies obtained from phase-contrast microscopy (Figure 5C) of a *C. difficile* 630 WT (*p*[ctl]), sRNA knock-out mutants (ΔSpoY/ΔSpoX-*p*[ctl]) and strains constitutively expressing the respective sRNA (ΔSpoY/ΔSpoX-*p*[ΔSpoY/ΔSpoX]). Samples (n=3 replicates) were taken at 12 h, 24 h and 48 h post inoculation of 70:30 liquid sporulation medium. ND: not determined – no viable spores. 2-way ANOVA with Dunnett's multiple comparison test was used to calculate statistical significance. Not significant (ns), $P \geq 0.05$; (*) $P$ 0.01 to 0.05; (****) $P < 0.0001$.

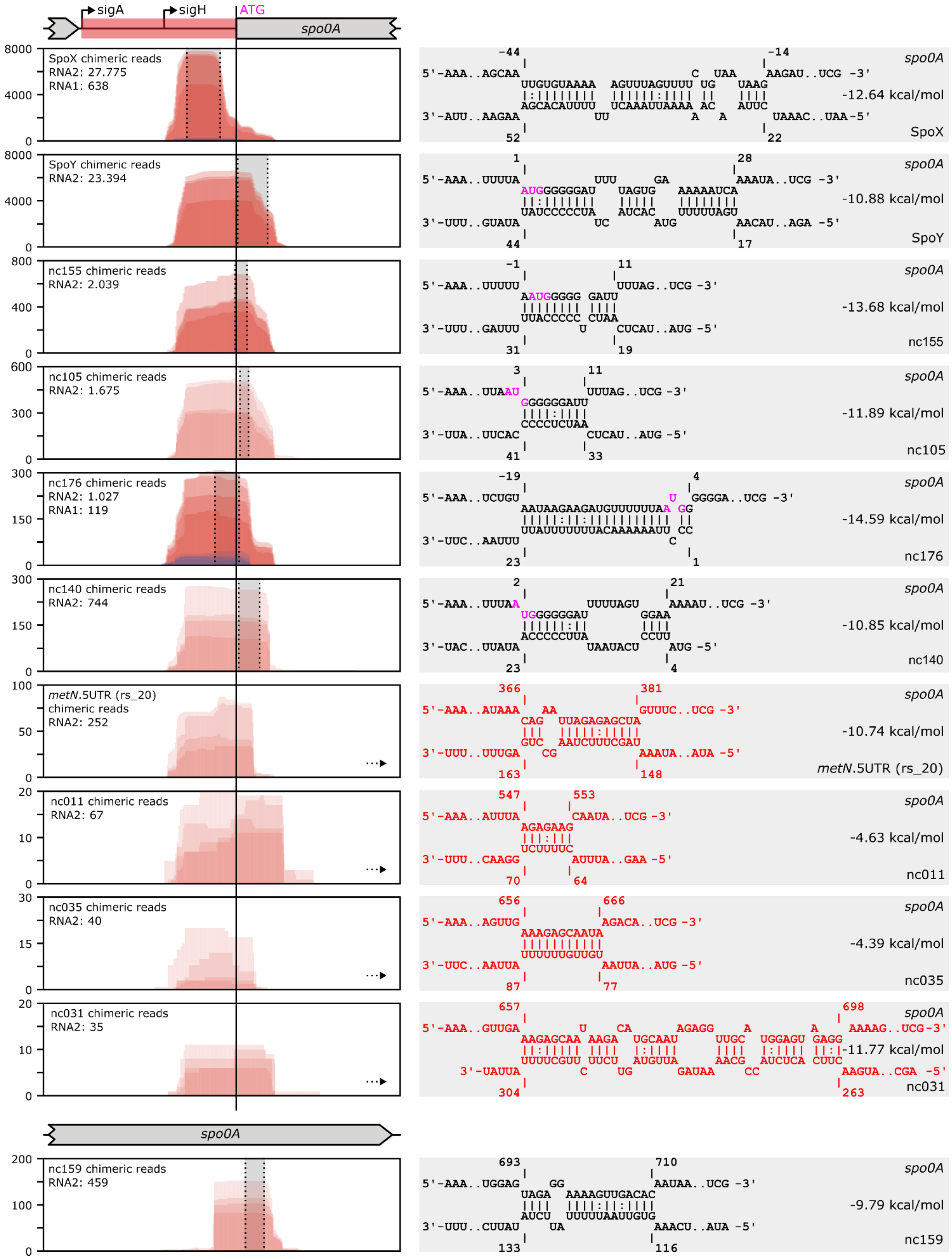

**Supplementary Figure 7: *spo0A* is a target of extensive sRNA-mediated post-transcriptional regulation.** On the left site, read coverage (y-axis) of *spo0A* by chimeric reads of all sRNA-*spo0A* interactions detected by RIL-seq analysis is depicted. The *spo0A* 5'UTR position including pTSS (sigA) and sTSS (sigH), start codon and coding sequence are marked on the x-axis. Chimeric reads covering *spo0A* were predominantly found at position 1 (RNA1) in a chimera and are color-coded in red. Chimeric reads found at position 2 (RNA2) are marked in blue. The number of chimeric reads covering each interaction is provided on the left. On the right site, base pairing information and location of the predicted binding sites (IntaRNA[2]) for each interaction are highlighted. *In silico* predictions that do not overlap with RIL-seq data are marked in red, and location of the interaction in relation to the RIL-seq peak is indicated by an arrow. Predicted interactions overlapping with RIL-seq data are shaded in gray and superimposed over the coverage plots. The *spo0A* nucleotide position is calculated relative to the *spo0A* start codon (highlighted in pink).

### SUPPLEMENTARY TABLES

#### Supplementary Table 1 (separate file)

**Number of fragments in RIL-seq sequencing libraries**. The table lists different libraries and corresponding statistics regarding the number of sequenced fragments. The following information is included:

- **Library** (name of the RIL-seq libraries)
- **Fragments** (number of raw fragments per library)
- **Remaining fragments after trimming and filtering** (number of remaining fragments after processing)
- **% remaining fragments after processing** (percentage of raw fragments remaining after trimming and filtering)
- **Fragments mapped in the first mapping step** (number of fragments mapped to CP010905.2 in the first mapping step of the RIL-seq computational pipeline)
- **% of mapped fragments (first mapping step)** (percentage of input fragments that were mapped in the first mapping step)
- **Fragments mapped in the second mapping step** (number of fragments that were mapped to CP010905.2 in the second mapping step of the RIL-seq computational pipeline)
- **Fragments mapped as singles** (number of fragments that were mapped to the same transcript or with less than 1000 nt distance)
- **Fragments mapped as chimeric** (number of fragments that were mapped to different transcripts or with more than 1000 nt distance)
- **% of mapped fragments (second mapping step)** (percentage of input reads that were mapped in the second mapping step)
- **% of single fragments** (percentage of mapped reads that were mapped as single fragments)
- **% of chimeric fragments** (percentage of mapped reads that were mapped as chimeric fragments)
- **S-chimeras** (number of S-chimeras in the output of the RIL-seq computational pipeline, no additional filtering)
- **% out of chimeric fragments** (percentage of fragments mapped as chimeric that were categorized as S-chimeras)
- **S-chimeras (S-chimeras with ≥25 interactions excluded)** (final number of S-chimeras in the unified data sets after manual curation and including the ≥25 interactions cut-off)

#### Supplementary Table 2 (separate file)

**RIL-seq RNA pairs identified in unified datasets of *C. difficile hfq*::3xFLAG strains.** The table includes all interactions between two RNAs, supported by statistically significant chimeras. Interactions supported by ≤25 chimeras are highlighted in red. Pairs can appear more than once if corresponding chimeric reads span multiple regions, such as an mRNA 5'UTR and CDS. Coordinates are based on the genome of *C. difficile* 630 (CP010905.2). The following information is included:

- **Start of RNA1 first read** (position of the first nucleotide of the most 5' chimera mapped to the first RNA)
- **Start of RNA1 last read** (position of the first nucleotide of the most 3' chimera mapped to the first RNA)
- **RNA1 strand** (the genome strand RNA1 was mapped to)
- **RNA1 match ID** (common locus tag of the match RNA1 mapped to)
- **RNA1 BioCyc ID** (BioCyc ID of RNA1)

- **RNA1 BioCyc Name** (BioCyc name of RNA1)
- **RNA1 match type** (type of RNA match, eg. sRNA, CDS, 3'UTR to which RNA1 maps)
- **sRNA annotation 1** (sRNA annotation if RNA1 maps to an sRNA)
- **RNA1 match start** (start of the match, RNA1 maps to, eg start position of a CDS)
- **RNA1 match end** (end of the match, RNA1 maps to, eg last position of a CDS)
- **RNA1 match strand** (the genome strand the match is located on)
- **Seq_RNA1** (peak sequence formed by RNA1 chimeric reads- the sequence is based on the position of RNA1 first read to start of RNA1 last read plus 50 nt)
- **Start of RNA2 first read** (position of the first nucleotide of the most 5' chimera mapped to the second RNA)
- **Start of RNA2 last read** (position of the first nucleotide of the most 3' chimera mapped to the second RNA)
- **RNA2 strand** (the genome strand RNA2 was mapped to)
- **RNA2 match ID** (common locus tag of the match RNA2 mapped to)
- **RNA2 BioCyc ID** (BioCyc ID of RNA2)
- **RNA2 BioCyc Name** (BioCyc name of RNA2)
- **RNA2 match type** (type of RNA match, eg. sRNA, CDS, 3'UTR to which RNA2 maps)
- **sRNA annotation 1** (sRNA annotation if RNA2 maps to an sRNA)
- **RNA2 match start** (start of the match, RNA2 maps to, eg start position of a CDS)
- **RNA2 match end** (end of the match, RNA2 maps to, eg last position of a CDS)
- **RNA2 match strand** (the genome strand the match is located on)
- **Seq_RNA2** (peak sequence formed by RNA2 chimeric reads- the sequence is based on the position of RNA2 first read to start of RNA2 last read plus 50 nt)
- **Interactions** (number of chimeric fragments supporting the interaction)
- **Other interactions of RNA1** (number of fragments in which the RNA1 appears at position one, including single fragments)
- **Other interactions of RNA2** (number of fragments in which the RNA2 appears at position 2, including single fragments)
- **Total other interactions** (number of fragments in the experiment excluding the above)
- **Odds ratio** ((K/L)/(M/N), where K= Number of chimeric fragments of RNA1-RNA2, L=number of other fragments involving RNA2, M=number of other fragments involving RNA1, N=number of all other fragments (that do not involve RNA1 and RNA2))
- **P-value (Fisher's exact test)** (p-value for observing at least this number of chimeric fragments given their background frequencies on Hfq. The Odds Ratio provides the effect size of the test)

**Supplementary Table 3 (separate file)**

**RIL-seq RNA pairs identified in unified datasets of *C. difficile* WT strains (control).** The table includes all interactions between two RNAs, supported by statistically significant chimeras. Interactions supported by ≤25 chimeras are highlighted in red. Pairs can appear more than once if corresponding chimeric reads span multiple regions, such as an mRNA 5'UTR and CDS. Coordinates are based on the genome of *C. difficile* 630 (CP010905.2). For table description, see Supplementary Table 2 above.

**Supplementary Table 4 (separate file)**

**RIL-seq analysis facilitates annotation of novel sRNAs**: The table lists potential novel sRNAs detected in the RIL-seq dataset. In addition to the RIL-seq information described above for Supplementary Table 2, the following information is given:

- **New sRNA** (name of new sRNA)
- **TSS** (approximate transcription start site of new sRNA, based on RIL-seq data and dRNA-seq/RIP-seq data published in Fuchs *et al.*[4])
- **PSS** (approximate processing site of new sRNA, based on RIL-seq data and dRNA-seq/RIP-seq data published in Fuchs *et al.*[4])
- **3'end** (approximate 3'end of new sRNA, based on RIL-seq data and dRNA-seq/RIP-seq data published in Fuchs *et al.*[4])
- **Length** (length of new sRNA, based on TSS, PSS and 3'end data)
- **Strand** (the genome strand the new sRNA is located on)
- **Type** (describes where the new sRNA is encoded, eg. 5'UTR of another gene)
- **Associated gene** (locus tag of the RNA with which the sRNA is associated with, eg. if the sRNA is encoded within the 5'UTR of another gene)
- **Chen *et al* (2011)** (if the sRNA was already annotated by Chen *et al.*[5], the corresponding name is given here)
- **Soutourina *et al* (2013)** (if the sRNA was already annotated by Soutourina *et al.*[6], the corresponding name is given here)
- **RIP-seq peak in Fuchs et al (2022)** (states if a RIP-seq peak corresponding to the sRNA was detected in the Hfq RIP-seq data published by Fuchs *et al.*[4])
- **Comment** (comments on location and peak profile of potential new sRNA)

**Supplementary Table 5 (separate file)**

**RIL-seq interactions overlapping with intra-operon RBS regions.** The table lists all RIL-seq interactions where either RNA1 or RNA2 overlaps with an intra-operon RBS region (25 nt upstream and 20 nt downstream of the respective start codon of the gene). Pairs can appear more than once if corresponding chimeric reads span multiple regions, such as an mRNA 5'UTR and CDS. Coordinates are based on the genome of *C. difficile* 630 (CP010905.2). In addition to the RIL-seq information described above for Supplementary Table 2, the following information is given:

- **RBS overlap RNA1** (states if RNA1 maps to an intra-operon RBS)
- **RBS overlap RNA2** (states if RNA2 maps to an intra-operon RBS)

**Supplementary Table 6**Plasmids and bacterial strains used in this study.

| Plasmid | Description | Origin |
|---|---|---|
| pJAK184 | To generate gene deletions or insertions in *C. difficile* by homologous recombination. Carrying *E. coli mazF* for counterselection in *C. difficile* | [4] |
| pFF-53 | Derived from pJAK184 for generating a *hfq*::3xFLAG strain | this study |

| | | |
|---|---|---|
| pSC-A-amp/kan | For cloning of PCR products using the StrataClone PCR Cloning Kit | Agilent Technologies, Inc. |
| pFF-162 | Derived from pSC-A-amp/kan for PCR amplification and subsequent *in vitro* transcription of SpoY (CDIF630nc_020) | this study |
| pFF-163 | Derived from pSC-A-amp/kan for PCR amplification and subsequent *in vitro* transcription of SpoY* (CDIF630nc_020 with a mutated seed region) | this study |
| pFF-164 | Derived from pSC-A-amp/kan for PCR amplification and subsequent *in vitro* transcription of SpoX (CDIF630nc_038, short isoform) | this study |
| pFF-245 | Derived from pSC-A-amp/kan for PCR amplification and subsequent *in vitro* transcription of SpoX* (CDIF630nc_038, short isoform with a mutated seed region) | this study |
| pFF-166 | Derived from pSC-A-amp/kan for PCR amplification and subsequent *in vitro* transcription of *spo0A* (CDIF630_01363 5'UTR starting from primary TSS plus 84 nt of CDS) | this study |
| pFF-248 | Derived from pSC-A-amp/kan for PCR amplification and subsequent *in vitro* transcription of of *spo0A**C* SpoY* (CDIF630_01363 5'UTR starting from primary TSS plus 84 nt of CDS with mutations compensating for the mutated SpoY seed region at the SpoY binding site) | this study |
| pFF-247 | Derived from pSC-A-amp/kan for PCR amplification and subsequent *in vitro* transcription of *spo0A**C* SpoX* (CDIF630_01363 5'UTR starting from primary TSS plus 84 nt of CDS with mutations compensating for the mutated SpoX seed region at the SpoX binding site) | this study |
| p JAK112 | To generate gene deletions in *C. difficile* 630 by homologous recombination. Carrying *E. coli codA* for counterselection in *C. difficile* | [4] |
| pFF-170 | Derived from pJAK112 for deletion of SpoY (CDIF630nc_020) | this study |
| pFF-171 | Derived from pJAK112 for deletion of SpoX (CDIF630nc_038) | this study |
| pDSW1728 | To monitor gene expression with a codon-optimized variant of mCherry (mCherryOpt) in *C. difficile*. Designed for cloning a promoter of interest upstream of *mCherryOpt* | [7] |

| | | |
|---|---|---|
| pFF-185 | *p*[*spo0A*] - Derived from pDSW1728 for constitutive expression of *spo0A* (CDIF630_01363 5'UTR starting from primary TSS plus 75 nt) of CDS fused to mCherryOpt, controlled by the *C. difficile* 630 *cwp2* promoter. | this study |
| pFF-186 | *p*[SpoY] - Derived from pDSW1728 for constitutive expression of SpoY (CDIF630nc_020), controlled by the *C. difficile* 630 *fdxA* promoter. | this study |
| pFF-191 | *p*[SpoY-*spo0A*] - Derived from pDSW1728 for constitutive co-expression of SpoY (CDIF630nc_020), controlled by the *C. difficile* 630 *fdxA* promoter, and *spo0A* (CDIF630_01363 5'UTR starting from primary TSS plus 75 nt of CDS) fused to mCherryOpt, controlled by the *C. difficile* 630 *cwp2* promoter. | this study |
| pFF-254 | *p*[SpoY*-*spo0A*] - Derived from pDSW1728 for constitutive co-expression of SpoY* (CDIF630nc_020 with a mutated seed region), controlled by the *C. difficile* 630 *fdxA* promoter, and *spo0A* (CDIF630_01363 5'UTR starting from primary TSS plus 75 nt) of CDS fused to mCherryOpt, controlled by the *C. difficile* 630 *cwp2* promoter. | this study |
| pFF-285 | *p*[SpoY*-*spo0A*$*^{C}$] - Derived from pDSW1728 for constitutive co-expression of SpoY* (CDIF630nc_020 with a mutated seed region), controlled by the *C. difficile* 630 *fdxA* promoter, and *spo0A* (CDIF630_01363 5'UTR starting from primary TSS plus 75 nt of CDS with mutations compensating for the mutated SpoY seed region at the SpoY binding site) fused to mCherryOpt, controlled by the *C. difficile* 630 *cwp2* promoter. | this study |
| pFF-187 | *p*[SpoX] - Derived from pDSW1728 for constitutive expression of SpoX (CDIF630nc_038, long isoform), controlled by the *C. difficile* 630 *fdxA* promoter. | this study |
| pFF-192 | *p*[SpoX-*spo0A*] - Derived from pDSW1728 for constitutive co-expression of SpoX (CDIF630nc_038, long isoform), controlled by the *C. difficile* 630 *fdxA* promoter, and *spo0A* (CDIF630_01363 5'UTR starting from primary TSS plus 75 nt of CDS) fused to mCherryOpt, controlled by the *C. difficile* 630 *cwp2* promoter. | this study |
| pFF-260 | *p*[SpoX*-*spo0A*] - Derived from pDSW1728 for constitutive co-expression of SpoX* (CDIF630nc_038, long isoform with a mutated seed region), controlled by the *C. difficile* 630 *fdxA* promoter, and *spo0A* (CDIF630_01363 5'UTR starting from primary TSS plus 75 nt) of CDS fused to mCherryOpt, controlled by the *C. difficile* 630 *cwp2* promoter. | this study |
| pFF-289 | *p*[SpoX*-*spo0A*$*^{C}$] - Derived from pDSW1728 for constitutive co-expression of SpoX* (CDIF630nc_038, long isoform with a mutated seed region), controlled by the *C. difficile* 630 *fdxA* promoter, and *spo0A* | this study |

| | | |
|---|---|---|
| | (CDIF630_01363 5'UTR starting from primary TSS plus 75 nt of CDS with mutations compensating for the mutated SpoX seed region at the SpoX binding site) fused to mCherryOpt, controlled by the *C. difficile* 630 *cwp2* promoter. | |
| pFF-207 | *p*[ctl] - Derived from pDSW1728, empty control vector | this study |

| **Strain** | **Relevant markers / Genotype** | **Origin** |
|---|---|---|
| *Escherichia coli* | | |
| TOP10 | F- mcrA Δ(mrr-hsdRMS-mcrBC) φ80lacZΔM15 ΔlacX74 nupG recA1 araD139 Δ(ara-leu)7697 galE15 galK16 rpsL(StrR) endA1 λ- | Invitrogen |
| CA434 | thi-1 hsdS20 (r-B, m-B) supE44 recAB ara-14 leuB5proA2 lacY1 galK rpsL20 (strR) xyl-5 mtl-1 | Dieter Jahn |
| FFS-204 | Top 10 carrying pFF-53 | this study |
| FFS-210 | CA434 carrying pFF-53 | this study |
| FFS-420 | StrataClone SoloPack Competent Cells carrying pFF-162 | this study |
| FFS-421 | StrataClone SoloPack Competent Cells carrying pFF-163 | this study |
| FFS-422 | StrataClone SoloPack Competent Cells carrying pFF-164 | this study |
| FFS-694 | StrataClone SoloPack Competent Cells carrying pFF-245 | this study |
| FFS-424 | StrataClone SoloPack Competent Cells carrying pFF-166 | this study |
| FFS-697 | StrataClone SoloPack Competent Cells carrying pFF-248 | this study |
| FFS-696 | StrataClone SoloPack Competent Cells carrying pFF-247 | this study |
| FFS-428 | Top 10 carrying pFF-170 | this study |
| FFS-450 | CA434 carrying pFF-170 | this study |
| FFS-429 | Top 10 carrying pFF-171 | this study |

| | | |
|---|---|---|
| FFS-451 | CA434 carrying pFF-171 | this study |
| FFS-479 | Top 10 carrying pFF-185 (*p*[*spo0A*]) | this study |
| FFS-480 | Top 10 carrying pFF-186 (*p*[SpoY]) | this study |
| FFS-505 | Top 10 carrying pFF-191 (*p*[SpoY-*spo0A*]) | this study |
| FFS-714 | Top 10 carrying pFF-254 (*p*[SpoY*-*spo0A*]) | this study |
| FFS-31 | Top 10 carrying pFF-285 (*p*[SpoY*-*spo0A*$^{*C}$]) | this study |
| FFS-481 | Top 10 carrying pFF-187 (*p*[SpoX]) | this study |
| FFS-506 | Top 10 carrying pFF-192 (*p*[SpoX-*spo0A*]) | this study |
| FFS-720 | Top 10 carrying pFF-260 (*p*[SpoX*-*spo0A*]) | this study |
| FFS-606 | Top 10 carrying pFF-289 (*p*[SpoX*-*spo0A*$^{*C}$]) | this study |
| FFS-502 | CA434 carrying pFF-185 (*p*[*spo0A*]) | this study |
| FFS-503 | CA434 carrying pFF-186 (*p*[SpoY]) | this study |
| FFS-529 | CA434 carrying pFF-191 (*p*[SpoY-*spo0A*]) | this study |
| FFS-753 | CA434 carrying pFF-254 (*p*[SpoY*-*spo0A*]) | this study |
| FFS-771 | CA434 carrying pFF-285 (*p*[SpoY*-*spo0A*$^{*C}$]) | this study |
| FFS-504 | CA434 carrying pFF-187 (*p*[SpoX]) | this study |
| FFS-530 | CA434 carrying pFF-192 (*p*[SpoX-*spo0A*]) | this study |
| FFS-759 | CA434 carrying pFF-260 (*p*[SpoX*-*spo0A*]) | this study |
| FFS-775 | CA434 carrying pFF-289 (p[SpoX*-spo0A*C]) | this study |
| FFS-564 | Top 10 carrying pFF-207 (*p*[ctl]) | this study |

| | | |
|---|---|---|
| FFS-586 | CA434 carrying pFF-207 (*p*[ctl]) | this study |
| *Clostridioides difficile* | | |
| 630 | 630 wild-type strain | DSMZ |
| FFS-220 | 630 *hfq*::3xFLAG | this study |
| FFS-491 | 630 ΔSpoY (ΔCDIF630nc_020) | this study |
| FFS-492 | 630 ΔSpoX (ΔCDIF630nc_038) | this study |
| FFS-536 | 630 ΔSpoY carrying pFF-185 (*p*[*spo0A*]) | this study |
| FFS-535 | 630 ΔSpoY carrying pFF-186 (*p*[SpoY]) | this study |
| FFS-537 | 630 ΔSpoY carrying pFF-191 (*p*[SpoY-*spo0A*]) | this study |
| FFS-779 | 630 ΔSpoY carrying pFF-254 (*p*[SpoY*-*spo0A*]) | this study |
| FFS-798 | 630 ΔSpoY carrying pFF-285 (*p*[SpoY*-spo0A*$^{C}$]) | this study |
| FFS-539 | 630 ΔSpoX carrying pFF-185 (*p*[*spo0A*]) | this study |
| FFS-538 | 630 ΔSpoX carrying pFF-187 (*p*[SpoX]) | this study |
| FFS-540 | 630 ΔSpoX carrying pFF-192 (*p*[SpoX-*spo0A*]) | this study |
| FFS-785 | 630 ΔSpoX carrying pFF-260 (*p*[SpoX*-*spo0A*]) | this study |
| FFS-802 | 630 ΔSpoX carrying pFF-289 (*p*[SpoX*-spo0A*$^{C}$]) | this study |
| FFS-591 | 630 WT carrying pFF-207 (*p*[ctl]) | this study |
| FFS-593 | 630 ΔSpoY carrying pFF-207 (*p*[ctl]) | this study |
| FFS-594 | 630 ΔSpoX carrying pFF-207 (*p*[ctl]) | this study |

**Supplementary Table 7:** DNA oligonucleotides used in this study.

| Oligo | Sequence (5'-3') | Purpose and Reference |
|---|---|---|
| *Plasmid construction* | | |
| FFO-364 | cgtagaaatacggtgtttttgttaccctaTTCTATGCAAATATATGAATATATGGATATTG | amplification of *hfq* CDS and upstream region for Gibson assembly into pJAK184 – for insertion of an *hfq* C-terminal 3XFLAG tag |
| FFO-365 | atggtctttgtagtcTCTGTTGTTATTATTATTGTTGTTTTG | amplification of *hfq* CDS and upstream region for Gibson assembly into pJAK184 – for insertion of an *hfq* C-terminal 3XFLAG tag |
| FFO-368 | gacgatgacaagtagATAATTAATTTAATTTAAGATGATTGAGAGG | amplification of *hfq* CDS and downstream region for Gibson assembly into pJAK184 – for insertion of an *hfq* C-terminal 3XFLAG tag |
| FFO-369 | gggattttggtcatgagattatcaaaaaggTACATAAGAATCGACTGGTGC | amplification of *hfq* CDS and downstream region for Gibson assembly into pJAK184 – for insertion of an *hfq* C-terminal 3XFLAG tag |
| FFO-366 | aataataacaacagaGACTACAAAGACCATGACGG | amplification of 3XFLAG tag for Gibson cloning into pJAK184– for insertion of an *hfq* C-terminal 3XFLAG tag |
| FFO-367 | aattaaattaattatCTACTTGTCATCGTCATCCTTG | amplification of 3XFLAG tag for Gibson cloning into pJAK184 – for insertion of an *hfq* C-terminal 3XFLAG tag |
| FFO-362 | CCTTTTTGATAATCTCATGACCAAAATC | linearization of pJAK184 for insertion of homology arms[8] |
| FFO-363 | TAGGGTAACAAAAAACACCGTATTTC | linearization of pJAK184 for insertion of homology arms[8] |
| FFO-958 | GTTTTTTTTAATACGACTCACTATAGGGagatagtagattacaatgatttttg | amplification of SpoY (CDIF630nc_020) for Strata cloning into pSC-A-amp/kan, adding a T7 promoter to the 5' end |

| | | |
|---|---|---|
| FFO-959 | aaaaaaaagagacagccc | amplification of SpoY (CDIF630nc_020) for Strata cloning into pSC-A-amp/kan, adding a T7 promoter to the 5' end |
| FFO-960 | AAAAAAAAGAGACAGCCCGTTTAAGAAGCT GTCATATATAATATGATAGTAG | amplification and mutation (seed region) of SpoY (CDIF630nc_020), for Strata cloning into pSC-A-amp/kan, adding a T7 promoter to the 5' end |
| FFO-961 | GTTTTTTTTAATACGACTCACTATAGGGaata taaaataacaaacaaatcttaacaaaaaattaaac | amplification of SpoX (CDIF630nc_038, short isoform) for Strata cloning into pSC-A-amp/kan, adding a T7 promoter to the 5' end |
| FFO-962 | aaaaataaagaaggcaacg | amplification of SpoX (CDIF630nc_038, short isoform) for Strata cloning into pSC-A-amp/kan, adding a T7 promoter to the 5' end |
| FFO-1261 | GTTTTTTTTAATACGACTCACTATAGGGAAT ATAAAATAACAAACAAATCTTAACAAAAAA TTTTTAAAAATTTAT | amplification and mutation (seed region) of SpoX (CDIF630nc_038, short isoform), for Strata cloning into pSC-A-amp/kan, adding a T7 promoter to the 5' end |
| FFO-1262 | AAAAATAAAGAAGGCAACGGGAAGCCTTCT TTCATATAAATTTTTAAAAA | amplification and mutation (seed region) of SpoX (CDIF630nc_038, short isoform), for Strata cloning into pSC-A-amp/kan, adding a T7 promoter to the 5' end |
| FFO-964 | GTTTTTTTTAATACGACTCACTATAGGGgagg cattaaaaaattttattattttatc | amplification of 5'UTR (pTSS) and start of CDS (84 nt) of *spo0A* for Strata cloning into pSC-A-amp/kan, adding a T7 promoter to the 5' end |
| FFO-966 | caaatactcttttaatacctgac | amplification of 5'UTR (pTSS) and start of CDS (84 nt) of *spo0A* for Strata cloning into pSC-A-amp/kan, adding a T7 promoter to the 5' end |
| FFO-1268 | taaaaatcatatcattaaaaaacatcttcttattacag | to insert compensatory mutations at the SpoY target site in *spo0A* (pTSS only) |
| FFO-1269 | ttttttaatgatatgatttttagtggaaaaaatcaaaatag | to insert compensatory mutations at the SpoY target site in *spo0A* (pTSS only) |

| | | |
|---|---|---|
| FFO-1259 | tttaaaaatatataattgcttcctaattatc | to insert compensatory mutations at the SpoX target site in *spo0A* (pTSS only) |
| FFO-1267 | atatatttttaaaagttttctgtaataagaag | to insert compensatory mutations at the SpoX target site in *spo0A* (pTSS only) |
| M13 rev | CAGGAAACAGCTATGAC | amplification of fragments inserted in pSC-A-amp/kan |
| M13 fwd | GTAAAACGACGGCCAGT | amplification of fragments inserted in pSC-A-amp/kan |
| FFO-977 | ACCCTAGAGCTCgagcatggtttaataaattagaaaatg | amplification of 1.2 kb homology arm upstream of SpoY (CDIF630nc_020) deletion region for insertion into pJAK112, SacI site |
| FFO-978 | ttaagaagctgtcattctactatctatatattattatacgatactactttttatatg | amplification of 1.2 kb homology arm upstream of SpoY (CDIF630nc_020) deletion region for insertion into pJAK112, SacI site |
| FFO-979 | tatatagatagtagaatgacagcttcttaaacggg | amplification of 1.2 kb homology arm downstream of SpoY (CDIF630nc_020) deletion region for insertion into pJAK112, BamHI site |
| FFO-980 | AAAAGGGGATCCaaagtatctatcaactctttatcaaaaag | amplification of 1.2 kb homology arm downstream of SpoY (CDIF630nc_020) deletion region for insertion into pJAK112, BamHI site |
| FFO-985 | ACCCTAGAGCTCtagtaagaggagacagagaaaaag | amplification of 1.2 kb homology arm upstream of SpoX (CDIF630nc_038) deletion region for insertion into pJAK112, SacI site |
| FFO-986 | gaccagttgtgcaaaataaaaaaataagctgtttctaaaatgatttc | amplification of 1.2 kb homology arm upstream of SpoX (CDIF630nc_038) deletion region for insertion into pJAK112, SacI site |
| FFO-987 | agcttatttttttattttgcacaactggtcattattaatg | amplification of 1.2 kb homology arm downstream of SpoX (CDIF630nc_038) |

| | | |
|---|---|---|
| | | deletion region for insertion into pJAK112, BamHI site |
| FFO-988 | AAAAGGGGATCCctctctattcatgcacaaaattg | amplification of 1.2 kb homology arm downstream of SpoX (CDIF630nc_038) deletion region for insertion into pJAK112, BamHI site |
| FFO-1004 | CATCAAGCTAGCaaaagtttatatcttttggttaattattacaataag | amplification of the *cwp2* promoter (80 nt upstream of TSS), inserting a NheI restriction site at the 5' end |
| FFO-1000 | attttttaatgcctcctcgagttaccaattataatatatatttgatattatttc | amplification of the *cwp2* promoter (80 nt upstream of TSS), inserting a XhoI restriction site at the 3' end |
| FFO-1001 | aattggtaactcgaggaggcattaaaaaattttattattttatcaattatc | amplification of 5'UTR (pTSS) and start of CDS (75 nt) of *spo0A*, inserting a XhoI restriction site at the 5' end |
| FFO-1002 | aatatcttcagatccaaatgctacatgttcgagctcttttaatacctgacaaaaatc | amplification of 5'UTR (pTSS) and start of CDS (75 nt) of *spo0A*, inserting a SacI restriction site at the 3' end |
| FFO-1056 | ttaaaagagctcgtatctaaaggagaagaagataatatg | amplification of mCherryOpt, inserting a SacI restriction site directly upstream of the second codon in the mCherryOpt CDS |
| FFO-1057 | cttataggatccttatttatataattcatccatacctcc | amplification of mCherryOpt |
| FFO-995 | catcaagctagcaacaagaatatcataataaagtttttgttg | amplification of the *fdxA* promoter (80 nt upstream of TSS), inserting a NheI restriction site at the 5' end |
| FFO-1005 | tgtaatctactatctcaataacattataacaaatattattgaatataacaattaaattaattc | amplification of the *fdxA* promoter (80 nt upstream of TSS), inserting a SpoY overlapping region at the 3'end |
| FFO-1006 | gttataatgttattgagatagtagattacaatgatttttgtac | amplification of SpoY (CDIF630nc_020), inserting a *fdxA* overlapping region at 5' end |
| FFO-1007 | CTTATAGGATCCaaaaaagacttctcatgagagaagcctttttctagaaaaaaaaagagacagcccg | amplification of SpoY (CDIF630nc_020), inserting a XbaI restriction site, *slpA* |

| | | |
|---|---|---|
| | | terminator and BamHI restriction site at 3' end |
| FFO-999 | tctagaaaaaaggcttctctcatgagaagtctttttaaaagtttatatcttttggttaattattacaataag | amplification of the *cwp2* promoter (80 nt upstream of TSS) and 5'UTR (pTSS) and start of CDS (75 nt) of *spo0A* fused to mCherryOpt from pFF-185, exchanging the NheI restriction site for a XbaI restriction site and *slpA* terminator at the 5' end |
| FFO-1057 | cttataggatccttatttatataattcatccatacctcc | amplification of the *cwp2* promoter (80 nt upstream of TSS) and 5'UTR (pTSS) and start of CDS (75 nt) of *spo0A* fused to mCherryOpt from pFF-185, including the BamHI restriction site at the 3' end |
| FFO-1008 | ttgttattttatattcaataacattataacaaatattattgaatataacaattaaattaattc | amplification of the *fdxA* promoter (80 nt upstream of TSS), inserting a SpoX overlapping region at the 3'end |
| FFO-1009 | gttataatgttattgaatataaaataacaaacaaatcttaacaaaaaattaaac | amplification of SpoX (CDIF630nc_038), inserting a *fdxA* overlapping region at 5' end |
| FFO-1010 | cttttttctagataaatatagaaagaactagcttaaaacataatataattac | amplification of SpoX (CDIF630nc_038, long isoform), inserting a XbaI restriction site at the 3' end |
| FFO-1264 | GTTATAATGTTATTGAATATAAAATAACAAACAAATCTTAACAAAAAATTTTTAAAAATTTAT | amplification and mutation (seed region) of SpoX (CDIF630nc_038), inserting a *fdxA* overlapping region at the 5' end |
| FFO-1263 | gccttctttattttataaaaaaataagctgtttctaaaatg | to mutate the SpoX seed region (CDIF630nc_038) |
| FFO-994 | CTTGTTgctagcttgatgcagaattc | to linearize pDSW1728 products, starting at the NheI restriction site |
| FFO-1205 | catcaagctagctataagttttaataaactttaaatagaaaaagg | to linearize pDSW1728 products downstream of the BamHI restriction site, inserting an additional NheI restriction site at the 5 'end |
| *Northern blot probes* | | |

| | | |
|---|---|---|
| FFO-942 | CAATTTTCAAAGGGTTAGGG | targeting CDIF630nc_152 |
| FFO-943 | AATCACCCAAACGCCAATAA | targeting CDIF630nc_153 |
| FFO-944 | AAATGGGGGAGATTGAGTAT | targeting CDIF630nc_155 |
| FFO-947 | AAAAAAGCACTCCCCCAGCA | targeting CDIF630nc_164 |
| FFO-948 | TTATAAGGAGTGCTTTGGTG | targeting CDIF630nc_165 |
| FFO-951 | GGCTCGATTCAGAAAATAT | targeting CDIF630nc_171 |
| FFO-317 | TAAGAAGCTGTCATATATAGGGGGA | targeting SpoY (CDIF630nc_020) |
| FFO-1014 | TTTGTTAAGATTTGTTTGTT | targeting SpoX (CDIF630nc_038) |
| FFO-352 | GAAACAGCCAAGTTATTCTA | targeting CDIF630nc_037 |
| CD76 | TCAGCGCTAGAGAGCTTAAC | targeting 5S rRNA[4] |
| *RT-qPCR primer* | | |
| FFO-1421 | ATGGGGGGATTTTTAGTGG | *spo0A*, 5' end[9] |
| FFO-1422 | TCATTTGAGTCTCTTGAACTGGTC | *spo0A*, 3' end[9] |
| FFO-1423 | GTTGGTTATGGCACTTGACAG | *sigE*, 5' end[9] |
| FFO-1424 | GACTGTGATATTCCAAGC | *sigE*, 3' end[9] |
| FFO-1425 | CAAGCAATTTAGGTCTAGTTAGGAGC | *sigF*, 5' end[9] |
| FFO-1426 | AAGCTTCACTTTCCATCTTTGCC | *sigF*, 3' end[9] |
| FFO-1427 | GGATAGAACAAGAGATGAGATACCC | *spoIVA*, 5' end[10] |
| FFO-1428 | CTGCTGCCTTTTCAAATGTC | *spoIVA*, 3' end[10] |
| FFO-1429 | GATGCTATCCCTACTGCAACG | *spoIIQ*, 5' end[10] |

| | | |
|---|---|---|
| FFO-1430 | GTCCTTCTGTTACCTTCTGTTC | *spoIIQ*, 3' end[10] |
| FFO-1431 | TGGTACAGAGGCTAACTATGTTCTTG | *sigK*, 5' end[11] |
| FFO-1432 | CTGGACAATTTCTCTTTCTCTAGG | *sigK*, 3' end[11] |
| FFO-1433 | GTGGTGTTAATACATCAGAACTTCC | *sigG*, 5' end[9] |
| FFO-1434 | GTTGAAAACCTTACATTTTGGC | *sigG*, 3' end[9] |
| FFO-1437 | ACAGAACAGTAGTACCAGG | *sspA*, 5' end[9] |
| FFO-1438 | CTATCTGTTGCTTTTTCCAGC | *sspA*, 3' end[9] |

**SUPPLEMENTARY REFERENCES**

1. Lorenz, R. *et al.* ViennaRNA package 2.0. *Algorithms Mol. Biol.* **6**, 26 (2011).

2. Mann, M., Wright, P. R. & Backofen, R. IntaRNA 2.0: enhanced and customizable prediction of RNA–RNA interactions. *Nucleic Acids Res.* **45**, W435–W439 (2017).

3. Bernhart, S. H. *et al.* Partition function and base pairing probabilities of RNA heterodimers. *Algorithms Mol. Biol.* **1**, 3 (2006).

4. Fuchs, M. *et al.* An RNA-centric global view of Clostridioides difficile reveals broad activity of Hfq in a clinically important gram-positive bacterium. *Proc. Natl. Acad. Sci.* **118**, (2021).

5. Chen, Y., Indurthi, D. C., Jones, S. W. & Papoutsakis, E. T. Small RNAs in the genus clostridium. *MBio* **2**, e00340-10 (2011).

6. Soutourina, O. A. *et al.* Genome-wide identification of regulatory RNAs in the human pathogen Clostridium difficile. *PLoS Genet.* **9**, e1003493 (2013).

7. Ransom, E. M., Ellermeier, C. D. & Weiss, D. S. Use of mCherry red fluorescent protein for studies of protein localization and gene expression in Clostridium difficile. *Appl. Environ. Microbiol.* **81**, 1652–1660 (2015).

8. Cartman, S. T., Kelly, M. L., Heeg, D., Heap, J. T. & Minton, N. P. Precise manipulation of the Clostridium difficile chromosome reveals a lack of association between the tcdC genotype and toxin p roduction. *Appl. Environ. Microbiol.* **78**, 4683–4690 (2012).

9. Oliveira, P. H. *et al.* Epigenomic characterization of Clostridioides difficile finds a conserved DNA methyltransferase that mediates sporulation and pathogenesis. *Nat. Microbiol.* **5**, 166–180 (2020).

10. Fimlaid, K. A. *et al.* Global analysis of the sporulation pathway of Clostridium difficile. *PLoS Genet.* **9**, e1003660 (2013).

11. Saujet, L. *et al.* Genome-wide analysis of cell type-specific gene transcription during spore formation in Clostridium difficile. *PLoS Genet.* **9**, e1003756 (2013).
